# Hepatic HIF2α modulates extra-hepatic disease-associated phenotypes during metabolic dysfunction-associated steatotic liver disease

**DOI:** 10.64898/2026.04.02.716074

**Authors:** Lorenz M.W. Holzner, Rosa M. Korpershoek, Youguo Niu, Anna Cochrane, Paula M. Darwin, Julia Babuta, Areesha Nazeer, Cecilia Castro, Alice P. Sowton, Alice E. Knapton, Benjamin D. Thackray, Julian L. Griffin, Zoe Hall, Dino A. Giussani, Rob C. I. Wüst, Andrew J. Murray

## Abstract

Metabolic dysfunction-associated steatotic liver disease (MASLD) afflicts more than one-third of adults globally, contributing significantly to an increased cardiovascular disease risk. Further, patients with severe liver disease experience muscle weakness (sarcopenic obesity) and fatigue. Hypoxia-inducible factor 2α (HIF2α) accumulates in the livers of MASLD patients and has been implicated in disease progression. Here we sought to understand the role of hepatic HIF2α in mediating hepatic and extra-hepatic features of MASLD. Using a well-validated obese mouse model of MASLD, we investigated the impact of hepatocyte-specific HIF2α deletion (hHIF2α^-/-^) on hepatic, cardiac and skeletal muscle metabolism, and cardiac function. Over 28 weeks, mice were exposed to a high-fat, high-fructose, high-cholesterol (GAN) diet, which induced obesity alongside hepatic steatosis, fibrosis and inflammation. In contrast to observations in lean mouse models of liver disease, hHIF2α^-/-^ did not protect against MASLD, despite greater hepatic NADH-supported mitochondrial respiration and higher intracellular sphingomyelin levels. Instead, in the hearts of GAN-fed mice, hHIF2α^-/-^ caused diacylglycerol accumulation independent of diet, accumulation of long-chain acyl-carnitines and exacerbation of ceramide accumulation. Langendorff-perfused hearts from hHIF2α^-/-^ mice showed systolic and diastolic dysfunction, including 24% lower left ventricular developed pressure and 34% lower maximal rate of relaxation (dP/dt_min_). However, isolated hearts from hHIF2α^-/-^ mice were protected against MASLD-associated sympathetic dominance, determined using autonomic receptor agonist stimulation. Both GAN-feeding and hHIF2α^-/-^ were associated with lower lean mass (14% and 5.4% lower than respective controls), whilst hHIF2α^-/-^ enhanced OXPHOS-associated protein levels in gastrocnemius muscle. Overall, hHIF2α^-/-^ resulted in detrimental extra-hepatic effects, including myocardial lipid accumulation, impaired cardiac function, and loss of whole-body lean mass, with no apparent protection against MASLD disease progression.

## Introduction

Metabolic dysfunction-associated steatotic liver disease (MASLD) is estimated to afflict 38% of the global adult population^1^. Prevalence is closely linked to the obesity epidemic, with body mass index (BMI) and metabolic abnormalities such as dyslipidaemia and type 2 diabetes mellitus being major risk factors^2^. MASLD is a spectrum disease, initially presenting as relatively benign fatty liver but with time progressing toward fibrosis and more severe metabolic dysfunction-associated steatohepatitis (MASH). MASH is characterised by hepatic inflammation, hepatocyte ballooning and death, and fibrosis. Eventually, cirrhosis or hepatocellular carcinoma can occur^3^. Currently, only the thyroid hormone receptor-beta agonist, resmetirom, has been specifically approved for MASLD treatment^4^, highlighting a need for improved understanding of pathophysiological mechanisms.

Studies in humans and rodents have highlighted a role for hepatic metabolism, with increased *de novo* lipogenesis^5,6^ and changes in mitochondrial fatty acid oxidation (FAO) associated with MASLD^7,8^. Resmetirom may itself regulate hepatic lipid metabolism^4^, further advocating for metabolic modulation as a therapeutic approach. One regulator of hepatic lipid metabolism is the hypoxia-inducible factor 2α (HIF2α)^9^. Commonly studied in the contexts of high-altitude adaptation^10–12^ and cancer^13^, HIF2α accumulates in the liver of patients with MASLD^14,15^ and regulates FAO, *de novo* lipogenesis and processes associated with disease progression^16^. Constitutive HIF activation, *via* hepatocyte-specific deletion of the Von Hippel-Lindau protein, suppressed FAO^9^ and increased expression of fatty acid synthesis genes in mice^17^, resulting in HIF2α-dependent onset of steatosis and inflammation. Further, a *EPAS1*/*HIF2A* mutation causing increased nuclear accumulation of HIF2α was associated with hepatic steatosis in humans^18^, and living at high altitude, i.e. in a lower oxygen environment, is associated with increased risk of developing MASLD^19^. In a lean mouse model of MASH (dietary choline deficiency), HIF2α deletion protected against disease progression^15^, but whether HIF2α contributes to MASLD in well-validated models of obesity that more closely resemble human pathology^20^ remains unclear.

MASLD is regarded as a multisystem disease owing to well-documented extra-hepatic manifestations^21^. Importantly, the main cause of death in MASLD patients is cardiovascular disease^22^, including cardiomyopathy^23^. Moreover, whilst lean mass is protective against MASLD development^24^, severe steatosis is associated with sarcopenia^25^ and fatigue is a common symptom in patients^26,27^. Metabolic alterations have long been implicated in skeletal muscle dysfunction^28^ and cardiomyopathy^29^, yet cardiac and skeletal muscle remain under-investigated in rodent models of MASLD. Critically, there is an urgent need to understand mechanisms linking pathological processes across organs, and investigation into tissue-specific and systemic metabolic alterations in the context of MASLD progression and hepatocyte-specific intervention could provide valuable insight into extrahepatic pathology.

Using a well-validated obese mouse model of MASLD, we investigated the effect of hepatocyte-specific deletion of HIF2α on liver pathology and lipid metabolism. Cardiac function, mitochondria and lipid metabolism were also studied, whilst the role of skeletal muscle was examined by measuring body composition and mitochondrial metabolism. Finally, the plasma lipidome was investigated as a potential mediator of crosstalk between the liver and other organs.

## Results

### Hepatocyte-specific deletion of HIF2α does not protect against GAN-induced metabolic dysfunction-associated steatotic liver disease

To determine whether hepatic HIF2α contributes to MASLD, wildtype mice and mice with a hepatocyte-specific deletion of HIF2α (hHIF2α^-/-^) were fed normal chow or the high-fat, high-fructose, high-cholesterol GAN diet for 28 weeks (Fig. 1A). Deletion of HIF2α was demonstrated through 79% lower expression of *Epas1* (encoding HIF2α) in livers of hHIF2α^-/-^ mice compared with wildtype mice (p < 0.0001). Notably, hepatic *Epas1* expression was two-fold higher in GAN-fed wildtype mice than chow-fed wildtype mice (p < 0.0001, Fig. 1B); aligning with findings in human patients and rodent models of MASLD^14,15^.

**Fig. 1:**
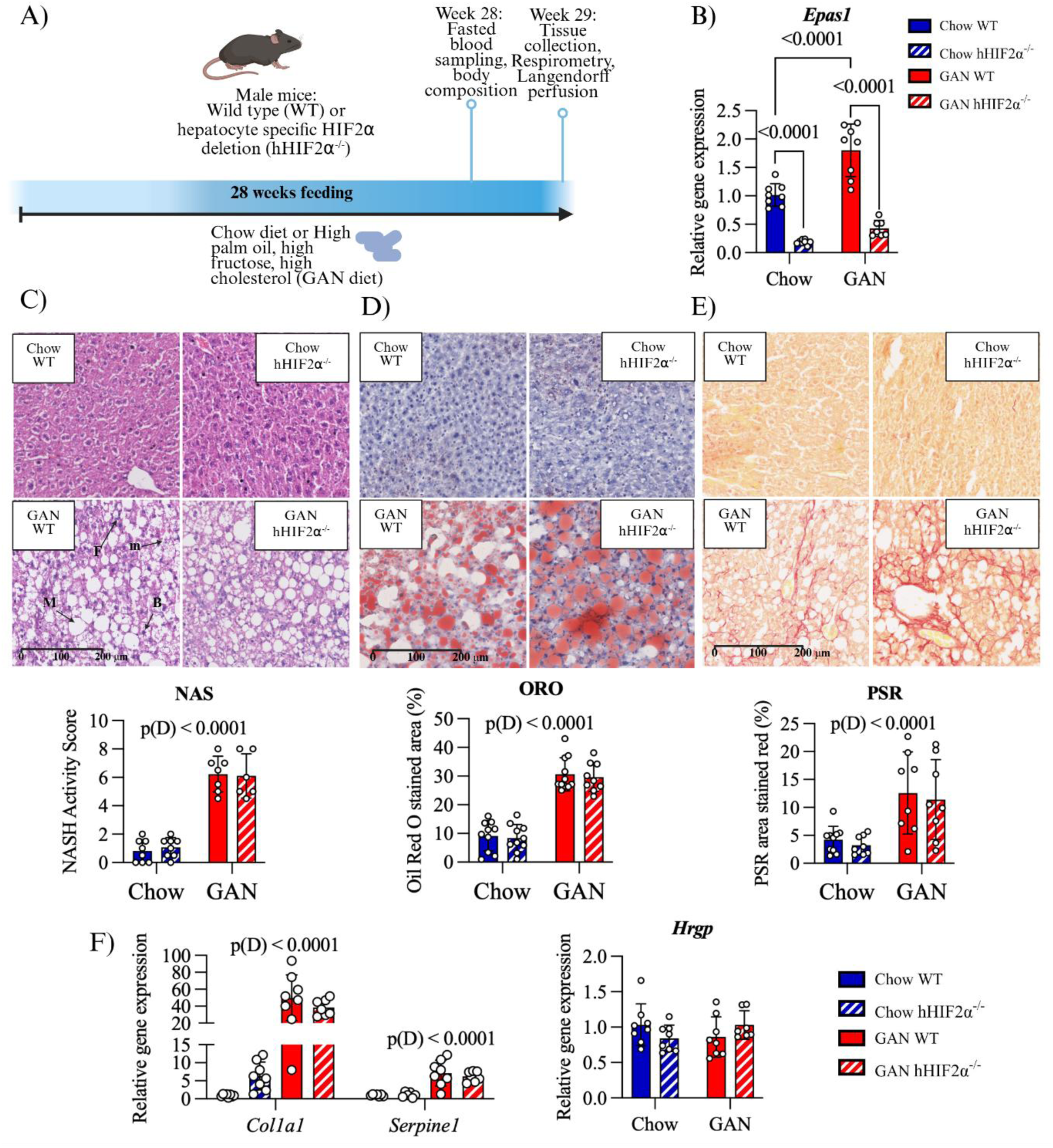
Hepatocyte-specific HIF2α deletion does not protect against metabolic dysfunction associated steatotic liver disease. A) Study design. Male mice (age 6-9 weeks, n = 13-14 per group) with hepatocyte-specific deletion of *Epas1*, the gene encoding HIF2α, (hHIF2α^-/-^), and wildtype (WT) littermates were fed either a chow control diet (Chow) or the high-fat, high-fructose, high-cholesterol GAN diet (GAN) for 28 weeks. Fasted blood sampling and body composition analysis (using tdNMR) was carried out after 14 and 28 weeks, with termination in week 29. Langendorff perfusion and high resolution respirometry experiments were carried out on fresh tissue. B) Relative gene expression of *Epas1* in livers from wildtype and hHIF2α^-/-^ chow and GAN-fed mice determined by RT-qPCR. n = 7-8 per group. C-D) Histological assessment of metabolic dysfunction-associated steatotic liver disease. C) Haematoxylin & Eosin stain, quantified as NASH activity score (NAS) ^89^. B – hepatocyte ballooning; F – inflammatory focus; M – macrovesicular steatosis; m – microvesicular steatosis. n = 6-8 per group. D) Oil red O (ORO) stain of neutral lipids, quantified as percentage area stained red below. n = 10-11 per group. E) Picrosirius red (PSR) stain of fibrosis, quantified as percentage area stained red below. n = 8-10 per group. F) Relative gene expression of genes involved in fibrosis (*Col1a1* and *Serpine1*) and inflammation (*Hrgp*). Data presented as mean ± SD Results of two-way ANOVA are shown on graphs. D = Diet main effect. In the case of a significant Diet x Genotype interaction, results of Tukey’s *post hoc* test are shown.

Body weights of GAN-fed mice were 32% higher than those of chow-fed mice (p < 0.0001), with no impact of genotype (Table 1). Adiposity was higher in GAN-fed mice than chow-fed mice, with a 3.2-fold greater fat mass measured by tdNMR (p < 0.0001) and 83% greater epididymal fat pad mass as a percentage of body mass (p < 0.0001). Lean mass, meanwhile, was 14% lower in GAN-fed mice than chow-fed mice (p < 0.0001). Further, hHIF2α^-/-^ mice had 5.4% lower lean mass (p = 0.0326) than wildtype mice.

**Table 1:**
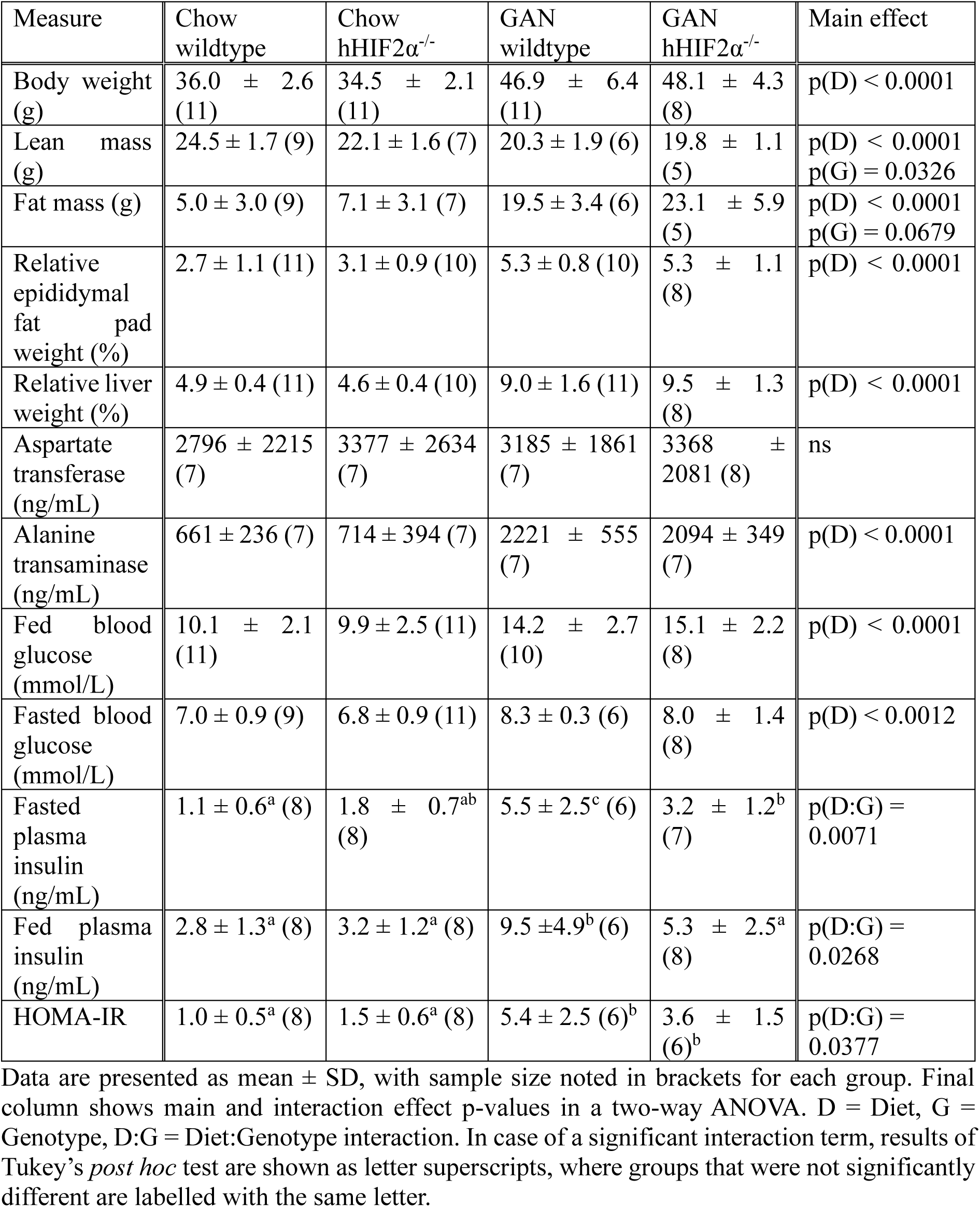
Body weight, body composition analysis, and plasma markers of liver function and glucose homeostasis in chow-fed and GAN-fed, wildtype and hHIF2α^-/-^ mice.

| Measure | Chow wildtype | Chow hHIF2α<sup>-/-</sup> | GAN wildtype | GAN hHIF2α<sup>-/-</sup> | Main effect |
| --- | --- | --- | --- | --- | --- |
| Body weight (g) | 36.0 ± 2.6 (11) | 34.5 ± 2.1 (11) | 46.9 ± 6.4 (11) | 48.1 ± 4.3 (8) | p(D) < 0.0001 |
| Lean mass (g) | 24.5 ± 1.7 (9) | 22.1 ± 1.6 (7) | 20.3 ± 1.9 (6) | 19.8 ± 1.1 (5) | p(D) < 0.0001<br>p(G) = 0.0326 |
| Fat mass (g) | 5.0 ± 3.0 (9) | 7.1 ± 3.1 (7) | 19.5 ± 3.4 (6) | 23.1 ± 5.9 (5) | p(D) < 0.0001<br>p(G) = 0.0679 |
| Relative epididymal fat pad weight (%) | 2.7 ± 1.1 (11) | 3.1 ± 0.9 (10) | 5.3 ± 0.8 (10) | 5.3 ± 1.1 (8) | p(D) < 0.0001 |
| Relative liver weight (%) | 4.9 ± 0.4 (11) | 4.6 ± 0.4 (10) | 9.0 ± 1.6 (11) | 9.5 ± 1.3 (8) | p(D) < 0.0001 |
| Aspartate transferase (ng/mL) | 2796 ± 2215 (7) | 3377 ± 2634 (7) | 3185 ± 1861 (7) | 3368 ± 2081 (8) | ns |
| Alanine transaminase (ng/mL) | 661 ± 236 (7) | 714 ± 394 (7) | 2221 ± 555 (7) | 2094 ± 349 (7) | p(D) < 0.0001 |
| Fed blood glucose (mmol/L) | 10.1 ± 2.1 (11) | 9.9 ± 2.5 (11) | 14.2 ± 2.7 (10) | 15.1 ± 2.2 (8) | p(D) < 0.0001 |
| Fasted blood glucose (mmol/L) | 7.0 ± 0.9 (9) | 6.8 ± 0.9 (11) | 8.3 ± 0.3 (6) | 8.0 ± 1.4 (8) | p(D) < 0.0012 |
| Fasted plasma insulin (ng/mL) | 1.1 ± 0.6<sup>a</sup> (8) | 1.8 ± 0.7<sup>ab</sup> (8) | 5.5 ± 2.5<sup>c</sup> (6) | 3.2 ± 1.2<sup>b</sup> (7) | p(D:G) = 0.0071 |
| Fed plasma insulin (ng/mL) | 2.8 ± 1.3<sup>a</sup> (8) | 3.2 ± 1.2<sup>a</sup> (8) | 9.5 ± 4.9<sup>b</sup> (6) | 5.3 ± 2.5<sup>a</sup> (8) | p(D:G) = 0.0268 |
| HOMA-IR | 1.0 ± 0.5<sup>a</sup> (8) | 1.5 ± 0.6<sup>a</sup> (8) | 5.4 ± 2.5 (6)<sup>b</sup> | 3.6 ± 1.5 (6)<sup>b</sup> | p(D:G) = 0.0377 |

Data are presented as mean ± SD, with sample size noted in brackets for each group. Final column shows main and interaction effect p-values in a two-way ANOVA. D = Diet, G = Genotype, D:G = Diet:Genotype interaction. In case of a significant interaction term, results of Tukey's *post hoc* test are shown as letter superscripts, where groups that were not significantly different are labelled with the same letter.

As expected, GAN feeding induced features of MASLD. NASH activity score, based on histology, was 6.7-fold higher in GAN-fed mice than chow-fed mice (p < 0.0001, Fig. 1C). Oil red O staining for steatosis was 3.4-fold higher in GAN-fed mice than chow-fed mice (p < 0.0001, Fig. 1D), whilst picrosirius red staining for fibrosis was 3.3-fold higher in GAN-fed mice (p < 0.0001, Fig. 1E). Fibrosis in GAN-fed mice was associated with increased expression of fibrosis markers *Col1a1* (induced 30-fold, p < 0.0001), and *Serpine1* (induced 6.3-fold, p < 0.0001, Fig 1F), with no significant effect of hHIF2α deletion on either marker. Gene expression of the proinflammatory marker histidine-rich glycoprotein (*Hrgp*) was not elevated in GAN-fed mice, and was unaffected by hHIF2α deletion.

In line with histological signs of pathology, liver weight as percentage of body weight was 1.9-fold greater in GAN-fed mice than chow-fed mice (p < 0.0001) and plasma levels of alanine transaminase were 3.1-fold higher in GAN-fed mice (p < 0.0001, Table 1). However, there was no apparent effect of hepatocyte HIF2α deletion on any MASLD-related feature. Thus, while HIF2α was expressed at higher levels in livers of wildtype GAN-fed mice than those of wildtype chow-fed mice, hHIF2α^-/-^ mice did not show protection against histological or plasma markers of MASLD, despite having a lower lean mass than wildtype mice.

### Hepatocyte-specific deletion of HIF2α prevents hyperinsulinaemia, but not hyperglycaemia

We further investigated whether hHIF2α^-/-^ mice had altered whole-body glucose homeostasis (Table 1). GAN feeding over 28 weeks resulted in elevated blood glucose, in both fed (43% higher than chow-fed mice, p < 0.0001) and fasted states (14% higher than chow-fed mice, p = 0.0012), whilst plasma insulin was higher in GAN-fed wildtype mice than chow-fed wildtype mice (fasted: 4.9-fold higher, p < 0.0001; fed: 3.4-fold higher, p < 0.001), resulting in a 5.4-fold greater HOMA-IR (a measure of insulin resistance) in GAN-fed wildtype mice than chow-fed wildtype mice (p < 0.0001). hHIF2α^-/-^ mice were not protected against hyperglycaemia, however GAN-fed hHIF2α^-/-^ mice had lower plasma insulin than GAN-fed wildtype mice in fed (42% lower, p = 0.0348) and fasted (44% lower, p = 0.0268) states. Therefore, HOMA-IR was also only 2.4-fold higher in GAN-fed hHIF2α^-/-^ mice than chow-fed hHIF2α^-/-^ mice (p = 0.0441).

### Deletion of hepatic HIF2α affects mitochondrial metabolism and gene expression in the liver, with limited impact on the hepatic lipidome

We had hypothesised that deletion of hepatocyte HIF2α would protect against MASLD through changes in mitochondrial respiration and lipid metabolism, and therefore assessed the expression of relevant genes using RT-qPCR (Fig. 2A). FAO-associated genes were expressed at higher levels in livers of GAN-fed mice than chow-fed mice: *Cpt1a* expression was 92% greater in GAN-fed wildtype mice than chow-fed wildtype mice (p = 0.0009), *Hadh* and *Pgc1a* were elevated in GAN-fed mice compared with chow-fed mice by 87% (p < 0.0001) and 72% (p < 0.0001), respectively, and *Ppara* was 87% greater in GAN-fed hHIF2α^-/-^ mice than chow-fed hHIF2^-/-^ mice (p = 0.0013). *Cpt1a* expression was 38% higher in GAN-fed hHIF2α^-/-^ mice than GAN-fed wildtype mice (p = 0.0327) but was unaffected by hHIF2α deletion in chow-fed mice. Hepatic expression of the lipogenic enzyme fatty acid synthase (*Fasn*) was 2.8-fold higher (p < 0.0001), and expression of *Srebf1*, which encodes SREBP1, a major regulator of hepatic *de novo* lipogenesis, was 43% higher in GAN-fed mice than chow-fed mice (p = 0.0025), though neither was affected by hHIF2α deletion. Thus, hHIF2α deletion increased the expression of FAO-associated genes, but did not alter *de novo* lipogenesis-associated gene expression.

**Fig. 2:**
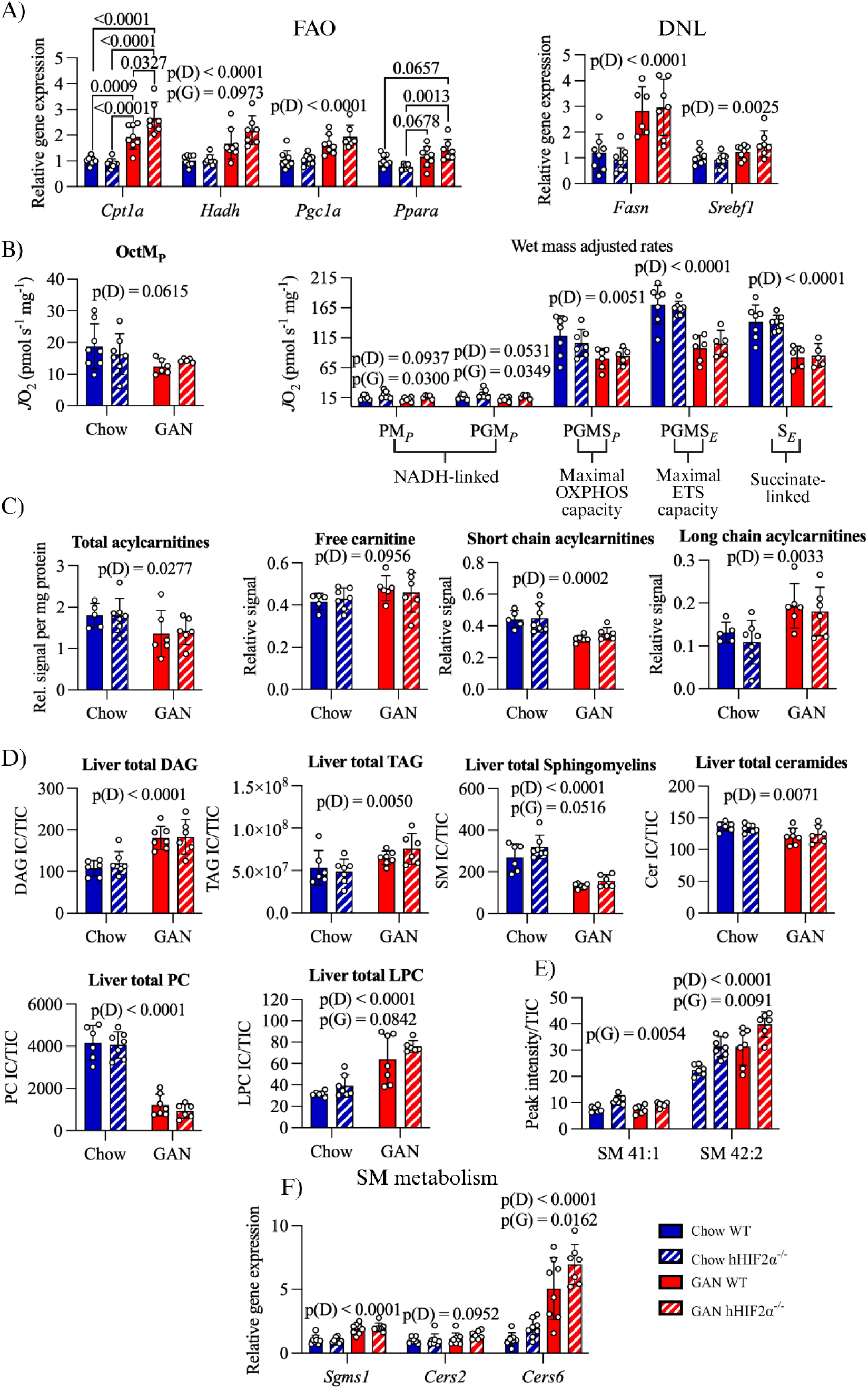
Hepatocyte-specific deletion of HIF2α is associated with changes in mitochondrial respiratory capacities and expression of fatty acid oxidation genes, but only limited lipidomic changes. A) Expression of fatty acid oxidation (FAO) and *de novo* lipogenesis (DNL) genes in livers of wildtype and hHIF2α^-/-^ chow and GAN-fed mice. n = 6-8 per group. B) Hepatic mitochondrial respiratory capacity assessed in a separate high resolution respirometry assay of liver homogenates from wildtype and hHIF2^-/-^ chow and GAN-fed mice. Rates adjusted to wet mass and corrected for residual oxygen consumption. Hepatic mitochondrial FAO capacity (OctM*_P_*) was measured using malate, octanoylcarnitine and ADP. n = 5-8 per group. In a separate assay, NADH-linked LEAK respiration (PM*_L_*) was measured using malate and pyruvate. Following addition of ADP, NADH-linked OXPHOS capacity (PM*_P_*, and PGM*_P_* with glutamate) was assessed. Maximal NADH- and succinate (S)-linked OXPHOS respiration (PGMS*_P_*) was stimulated using succinate. Following titration of the uncoupler FCCP electron transfer system (ETS) capacity (PGMS*_E_*) was determined. Inhibition of complex I using rotenone was used to limit electron flow to complex II (S*_E_*). n = 5-7 per group. C) Hepatic acylcarnitine species measured using liquid chromatography – mass spectrometry. Total acylcarnitine levels normalised to mass of protein pellet after extraction. Levels of free carnitine, and short and long chain acylcarnitines expressed relative to total acylcarnitine. n = 5-7 per group. D) Hepatic lipidome of wildtype and hHIF2α^-/-^ chow and GAN-fed mice assessed using liquid chromatography – mass spectrometry. Signals belonging to all lipids of a class were summed and normalised to total signal (total ion count, TIC). DAG – diacylglycerol; TAG – triacylglycerol; PC – phosphatidylcholine; LPC – lysophosphatidylcholine. n = 6-7 per group. E) Lipids significantly affected by hepatic HIF2α deletion using two-way ANOVA adjusted for multiple comparisons. SM – sphingomyelin. n = 6-7 per group. F) Expression of sphingomyelin metabolism genes. n = 6-8 per group Data presented as mean ± SD. Results of two-way ANOVA are shown on graphs. D = Diet, G = Genotype main effect. In the case of a significant Diet x Genotype interaction, results of Tukey’s *post hoc* test are shown.

Next, we investigated whether the changes in gene expression induced changes in FAO. High-resolution respirometry (Fig. 2B) revealed a trend towards lower hepatic oxidative phosphorylation capacity supported by octanoylcarnitine (medium chain acylcarnitine bypassing CPT1) and malate (OctM*_P_*) in GAN-fed compared with chow-fed mice (Fig 2B, p = 0.0615), with no effect of hepatic HIF2α deletion. The hepatic acylcarnitine pool was analysed for further insight (Fig. 2C). Total acylcarnitine levels were 23% lower in GAN-fed mice than chow-fed mice (p = 0.0277) but not affected by genotype. There were 25% lower relative levels of short-chain (p = 0.0002) and 56% higher relative levels of long-chain acylcarnitines (p = 0.0033) in GAN-fed compared with chow-fed mice. Medium-chain acylcarnitines were not affected by diet (Supplementary Fig. S1A). Collectively, these data suggest relative hepatic FAO insufficiency in GAN-fed mice compared with chow-fed mice, with no effect of hepatic HIF2α deletion.

High-resolution respirometry revealed further GAN diet-induced changes, including 32% lower maximal oxidative phosphorylation capacity (p = 0.0051), 39% lower maximal electron transfer capacity (p < 0.0001), and 40% lower succinate (complex II)-linked respiration (p < 0.0001). Hepatocyte-specific HIF2α deletion was only associated with changes in NADH-linked respiratory capacities, which were 31% greater in hHIF2α^-/-^ mice than wildtype mice (p = 0.0300 and 0.0349), with similar effects seen after normalisation to maximal electron transfer capacity (Supplementary Fig. 1B). These changes were not associated with altered expression of the complex I subunit *Ndufa9* (Supplementary Fig. S1C).

Investigation of the wider hepatic lipidome revealed clear diet-associated differences, but again only limited effects of hepatocyte HIF2α deletion. Analysis of relative levels of different lipid classes (Fig. 2D) revealed that diacylglycerol levels were 60% higher (p < 0.0001), triacylglycerols 38% higher (p = 0.0050), sphingomyelins 51% lower (p < 0.0001), ceramides 9% lower (p = 0.0071), phosphatidylcholines 74% lower (p < 0.0001) and lysophosphatidylcholines 99% higher (p < 0.0001) in GAN-fed compared with chow-fed mice. No lipid class was significantly affected by hepatic HIF2α deletion. Principal component analysis similarly highlighted separation by diet but not genotype (Supplementary Fig. S1E). Therefore, GAN diet-feeding resulted in lower hepatic mitochondrial respiratory capacity, and a shift in the lipidome towards glycerolipids, whilst hepatic HIF2α deletion was associated with greater NADH-linked oxidative phosphorylation capacity.

While analysis of lipid classes and multivariate approaches did not reveal any significant effect of hepatocyte-specific HIF2α deletion, analysis of individual lipids by two-way ANOVA (adjusted for multiple comparisons, Fig. 2E) revealed two sphingomyelin (SM) species that were present at higher relative levels in livers of hHIF2α^-/-^ mice. SM 41:1 was 34% higher (p = 0.0054) and SM 42:2 14% higher (p = 0.0091) in hHIF2α^-/-^ mice than wildtype mice, with the latter also being 22% higher in GAN-fed mice than chow-fed mice (p < 0.0001). Expression of *Sgms1*, encoding a sphingomyelin synthase, and *Cers2*, encoding ceramide synthase 2 (responsible for longer chain ceramides, which are converted into sphingomyelins) were unaffected by genotype, though *Sgms1* expression was 93% higher in GAN-fed mice than chow-fed mice (p < 0.0001). *Cers6*, which synthesises palmitate-containing ceramides, was expressed at 4-fold higher levels in GAN-fed than chow-fed mice (p < 0.0001) and was 58% higher in hHIF2α^-/-^ mice than wildtype mice (p = 0.0162), highlighting a specific effect of hHIF2α deletion on hepatic sphingolipid metabolism.

### Hepatocyte-specific HIF2α is associated with basal cardiac dysfunction but protects against GAN-induced sympathetic dominance

As MASLD has been linked with cardiovascular disease risk, we investigated basal and stimulated function in chow and GAN-fed wildtype and hHIF2α^-/-^ mouse hearts *ex vivo,* using the Langendorff-perfused heart preparation. Basal systolic function (Fig. 3A) was impacted by both diet and genotype: left ventricular developed pressure (LVDP) being 25% lower in GAN-fed mice than chow-fed mice (p = 0.0278) and 24% lower in hHIF2α^-/-^ mice than wildtype mice (p = 0.0384). Similarly, the maximal rate of contraction, dP/dt_max_, was 32% lower in hHIF2α^-/-^mice than wildtype mice (p = 0.0035). Basal diastolic function was also affected by both diet and genotype (Fig. 3B). While left ventricular end-diastolic pressure (LVEDP) was not altered, the maximal rate of relaxation, dp/dt_min_, was 34% lower in hHIF2α^-/-^ mice than wildtype mice (p = 0.0007). The relaxation time constant (τ) was 57% higher in GAN-fed mice than chow-fed mice (p = 0.0063).

**Fig. 3:**
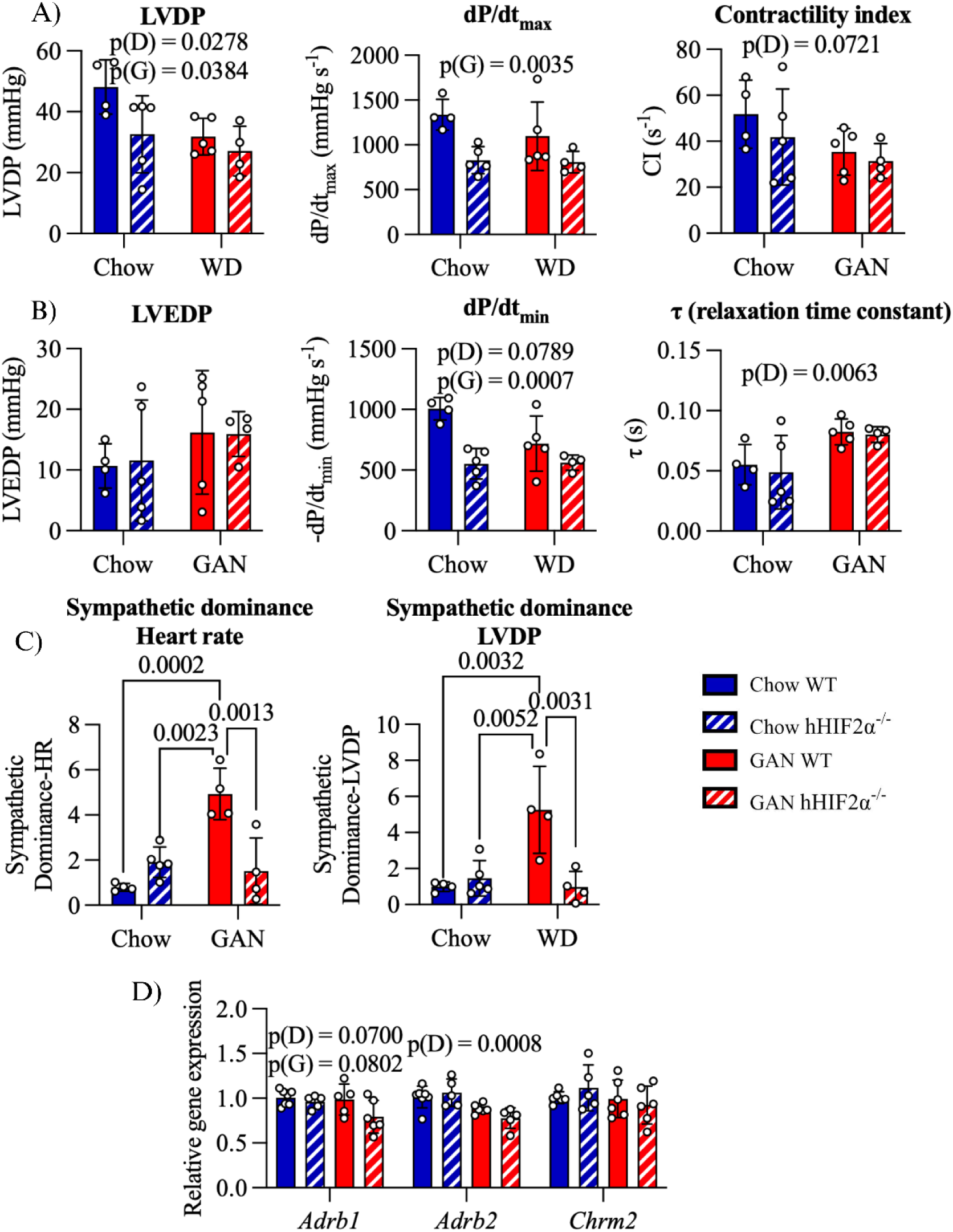
Hepatocyte-specific deletion of HIF2α and GAN-induced MASLD are associated with cardiac dysfunction but hepatic HIF2α deletion protects against GAN-induced cardiac sympathetic dominance. Cardiac left ventricular function in hHIF2α^-/-^ and wildtype chow and GAN-fed mice assessed in Langendorff perfused hearts. n = 4-5 per group. A) Systolic function: Left ventricular developed pressure (LVDP), maximal rate of pressure change dP/dt_max_, and the contractility index. B) Diastolic function: Left ventricular end-diastolic pressure (LVEDP), minimal rate of pressure change dP/dt_min_, and the relaxation time constant τ. C) Sympathetic dominance ratio of heart rate and LVDP. Chronotropic and inotropic responsiveness to muscarinic acetylcholine receptor stimulation using carbachol, and β-adrenergic receptor stimulation using isoprenaline, and the ratio of the response to the maximal isoprenaline dose relative to the maximal carbachol dose (sympathetic dominance ratio). A higher value denotes relative sympathetic dominance. D) Cardiac expression of β_1_- (*Adrb1*), β_2_-adrenergic (*Adrb2*) and M_2_-muscarinic acid receptor (*Chrm2*). n = 5-7 per group. Data presented as mean ± SD. Results of two-way ANOVA are shown on graphs. D = Diet, G = Genotype main effect. In the case of a significant Diet x Genotype interaction, results of Tukey’s *post hoc* test are shown.

In addition to basal function, we investigated cardiac responsiveness to autonomic stimulation, using agonists of β-adrenergic (isoprenaline) and muscarinic cholinergic (carbachol) receptors. In wildtype mice, GAN-feeding recapitulated the cardiac sympathetic dominance seen in MASLD^30^. The ratio of the maximal chronotropic response to isoprenaline and carbachol, an index of sympathetic dominance^31^ (Fig. 3C), was 6.2-fold higher in GAN-fed wildtype mouse hearts than those of chow-fed wildtype mice (p = 0.0002), but also 3.3-fold higher in GAN-fed wildtype mouse hearts than those of GAN-fed hHIF2α^-/-^ mice (p = 0.0013). Similarly, the inotropic sympathetic dominance ratio was 5.4-fold higher in GAN-fed wildtype mouse hearts than chow-fed wildtype mouse hearts (p = 0.0032), and 5.4-fold higher in GAN-fed wildtype mice than GAN-fed hHIF2α^-/-^ mice (p = 0.0031). In both cases, this was associated with a greater difference in carbachol than isoprenaline sensitivity (Supplementary Fig. S2). Overall, hepatocyte-specific deletion of HIF2α was associated with systolic and diastolic dysfunction in the heart, but protection against GAN-induced cardiac sympathetic dominance.

We next measured cardiac expression of β-adrenergic and muscarinic receptors (Fig. 3D) to determine whether changes in sympathetic dominance might be attributed to altered receptor expression. There were no significant differences in the expression of any measured receptors between GAN-fed wildtype and hHIF2α^-/-^ mice, although there was a trend towards a genotype (p = 0.0802) and a diet effect (p = 0.0700) on *Adrb1* (encoding the β_1_-adrenergic receptor) expression and *Adrb2* (β_2_-adrenergic receptor), which couples with the Gi protein ^32^, showed lower expression in GAN-fed mice than chow-fed mice (p = 0.0008). *Chrm2* (muscarinic receptor 2) expression was unchanged. Thus, hHIF2α deletion-mediated protection against sympathetic dominance was not associated with altered autonomic receptor expression.

### Hepatocyte-specific deletion of HIF2α is associated with accumulation of putatively lipotoxic and signalling lipids in the heart

To determine whether metabolic changes could underlie cardiac phenotypes of hHIF2α^-/-^ and GAN-fed mice, we investigated cardiac lipids. Histology revealed a 7.2-fold higher accumulation of neutral lipids in the hearts of GAN-fed mice than chow-fed counterparts (p = 0.0003, Fig 4A). Similarly, cardiac lipidomic analysis (Fig. 4B) showed 38% higher relative triacylglycerol levels in GAN-fed mice than chow-fed mice (p = 0.0008). Relative sphingomyelin and phosphatidylcholine levels were lower in GAN-fed than chow-fed mice, by 21% (p < 0.0001) and 14% (p = 0.0005), respectively. Lipid classes associated with lipotoxicity were elevated in the hearts of hHIFα^-/-^ mice. Diacylglycerol levels were 13% higher in hHIF2α^-/-^ mice than wildtype mice (p = 0.0087), while ceramide levels were 45% greater in GAN-fed wildtype than chow-fed wildtype mice (p < 0.0103) and a further 25% greater in the GAN-fed hHIF2α^-/-^ mouse hearts than those of GAN-fed wildtype mice (p = 0.0430), with no difference between chow-fed groups. Further, lysophosphatidylcholines, which have signalling roles, were 21% higher in hHIF2α^-/-^ mice than wildtype mice (p = 0.0146), with the ratio of lysophosphatidylcholine-to-phosphatidylcholine being 39-54% higher in GAN-fed hHIF2α^-/-^mice than all other groups.

**Fig. 4:**
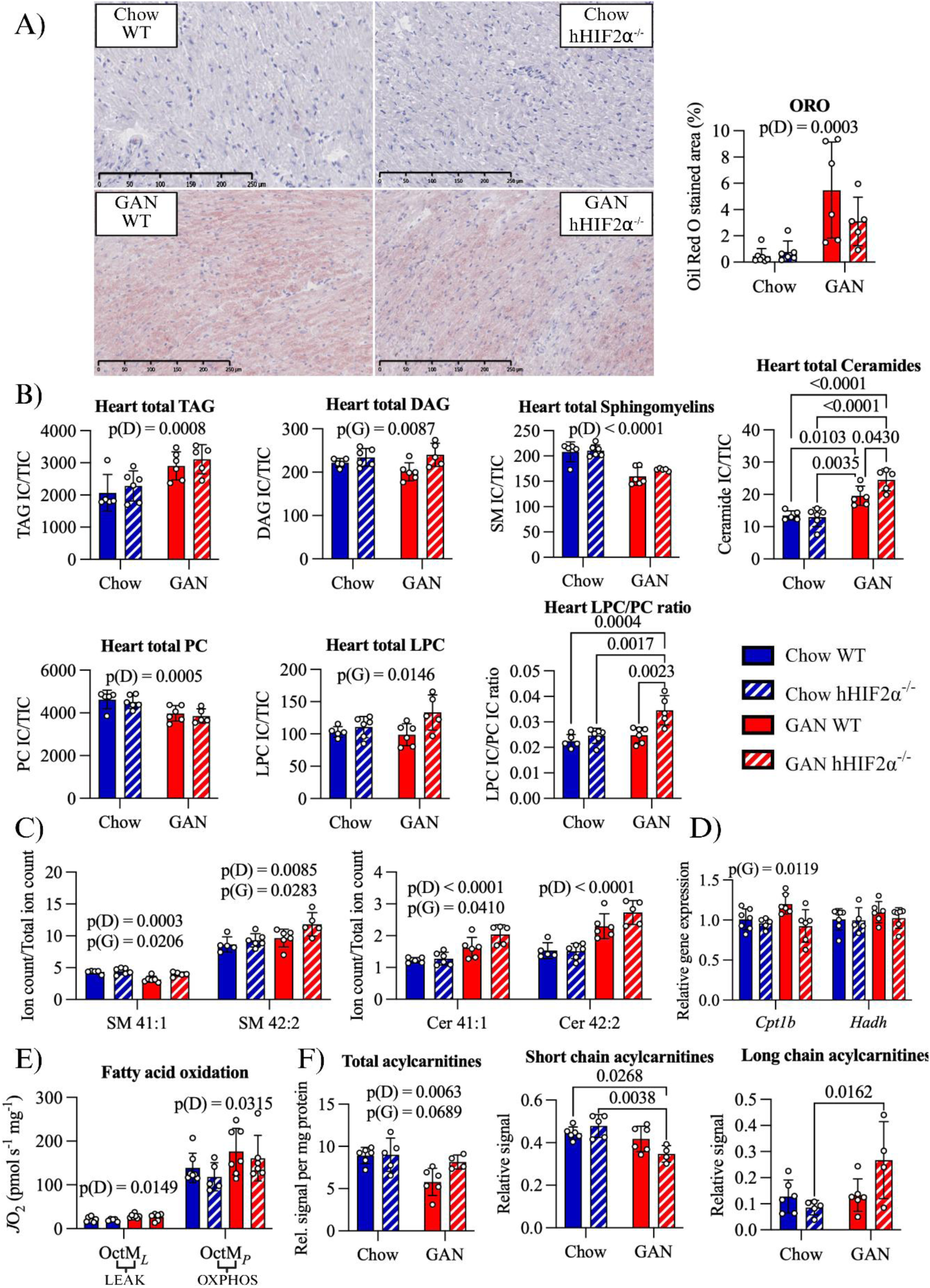
Hepatocyte-specific deletion of HIF2α is associated with cardiac accumulation of putatively lipotoxic and signalling lipids, particularly in the context of MASLD. A) Cardiac neutral lipid accumulation as measured using ORO staining. Example images and quantification of stained area. n = 5-7 per group. B) Sums of signals for lipid classes relative to total signal of the cardiac lipidome. TAG – triacylglycerol, DAG – diacylglycerol, PC – phosphatidylcholine, LPC – lysophosphatidylcholine. n = 5-6 per group. C) Relative cardiac levels of sphingomyelins SM 41:1 and 42:2, which were affected by hHIF2α deletion in the liver, and their corresponding ceramides, normalised to total signal. n = 5-6 per group. D) Expression of FAO genes *Cpt1b* and *Hadh*. n = 5-7 per group. E) Cardiac fatty acid oxidation capacity measured using high resolution respirometry in permeabilised cardiac fibres from wildtype and hHIF2^-/-^ chow and GAN-fed mice. Rates adjusted to wet mass and corrected for residual oxygen consumption. FAO (F-pathway) supported LEAK respiration (OctM*_L_*) was measured using octanoylcarnitine and malate. Following addition of ADP, F-pathway linked OXPHOS capacity (OctM*_P_*) was assessed. F) Cardiac acylcarnitine species measured using liquid chromatography – mass spectrometry. Total acylcarnitine levels, and relative levels of short and long-chain acylcarnitines to total acylcarnitine are shown. n = 4-6 per group. Data presented as mean ± SD. Results of two-way ANOVA are shown on graphs. D = Diet, G = Genotype main effect. In the case of a significant Diet x Genotype interaction, results of Tukey’s *post hoc* test are shown.

We also specifically analysed the two sphingomyelin species elevated in the livers of hHIF2α^-/-^ mice, as well as their corresponding ceramides, using two-way ANOVA (Fig. 4C). SM 41:1 was 18% lower in the hearts of GAN-fed mice than chow-fed mice (p = 0.0003), and 14% higher in the hearts of hHIF2α^-/-^ mice than wildtype mice (p = 0.0206). SM 42:2 was 19% higher in the hearts of GAN-fed mice than chow-fed mice (p = 0.0085) and 15% higher in the hearts of hHIF2α^-/-^ mice than wildtype mice (p = 0.0283). Cer 41:1 was 46% higher in hearts of GAN-fed mice than those of chow-fed mice (p < 0.0001) and 15% higher in hHIF2α^-/-^ mice than wildtype mice (p = 0.0410), while Cer 42:2 was affected only by diet, being 50% higher in GAN-fed mice than chow-fed mice (p < 0.0001). These sphingomyelins, and the related ceramide 41:1, were thus similarly altered in the hearts and liver of hHIF2α^-/-^ mice, suggesting a possible mechanism for inter-organ signalling.

Expression of *Cpt1b* (Fig. 4D), the rate-limiting enzyme of long-chain FAO, was 14% lower in hHIF2α^-/-^ mice than wildtype mice (p = 0.0119), while *Hadh* expression was not different between groups. High-resolution respirometry (Fig. 4E) did not reveal any genotype-related changes in cardiac mitochondrial function, though GAN-fed mice had 33% greater oxidative phosphorylation capacities supported by octanoylcarnitine and malate (p = 0.0315). Total acylcarnitine levels were lower in GAN fed mice (p = 0.0063) and trended higher in hHIF2α^-/-^ mice (p = 0.0689). Relative levels of short-chain acylcarnitines were 25% lower in GAN-fed hHIF2α^-/-^ mice than in chow-fed wildtype mice (p = 0.0268), and relative levels of long-chain acylcarnitines were 3.1-fold higher in GAN-fed than chow-fed hHIF2α^-/-^ mice (p = 0.0162, Fig. 4F). These data suggest relative insufficiency in cardiac FAO in GAN-fed hHIF2α^-/-^ mice compared with wildtype mice.

Therefore, neutral lipids accumulated in the hearts of GAN-fed mice, while lipotoxic and signalling lipids, including species altered in the liver, were found at higher levels in the hearts of hHIF2α^-/-^ mice, which also had lower expression of *Cpt1b* and, in the context of GAN-feeding, accumulation of long-chain acylcarnitines.

### Mice lacking hepatic HIF2α exhibit changes in skeletal muscle oxidative phosphorylation gene expression

We next investigated the skeletal muscle phenotype of GAN-fed and hHIF2α^-/-^ mice, both of which had lower whole-body lean mass. In the mixed fibre-type gastrocnemius, relative to wet weight, respiratory capacity (measured in freeze-thawed samples) was unchanged (see Supplementary Fig. S3A), although there was a trend towards greater NADH-linked oxidative phosphorylation relative to maximal capacity (p = 0.0837, Fig. 5A) in hHIF2α^-/-^ mice than wildtype mice. Citrate synthase activity, a marker of mitochondrial content, tended to be lower (p = 0.0518, Fig. 5B) in hHIF2α^-/-^ mice compared with wildtype mice. Protein levels of subunits of mitochondrial complex I (p = 0.0208) and II (p = 0.0455) were, respectively, 1.8- and 2-fold higher in hHIF2α^-/-^ mice than wildtype mice, and there was a trend (p = 0.0780) towards higher levels of the complex III subunit UQRC2 in hHIF2α^-/-^ than wildtype mice (Fig. 5C). Measured subunits of complex IV and ATP synthase were not different between groups. These results indicate an altered distribution of the proteins related to oxidative phosphorylation in skeletal muscle in hHIF2α^-/-^ mice.

**Fig. 5:**
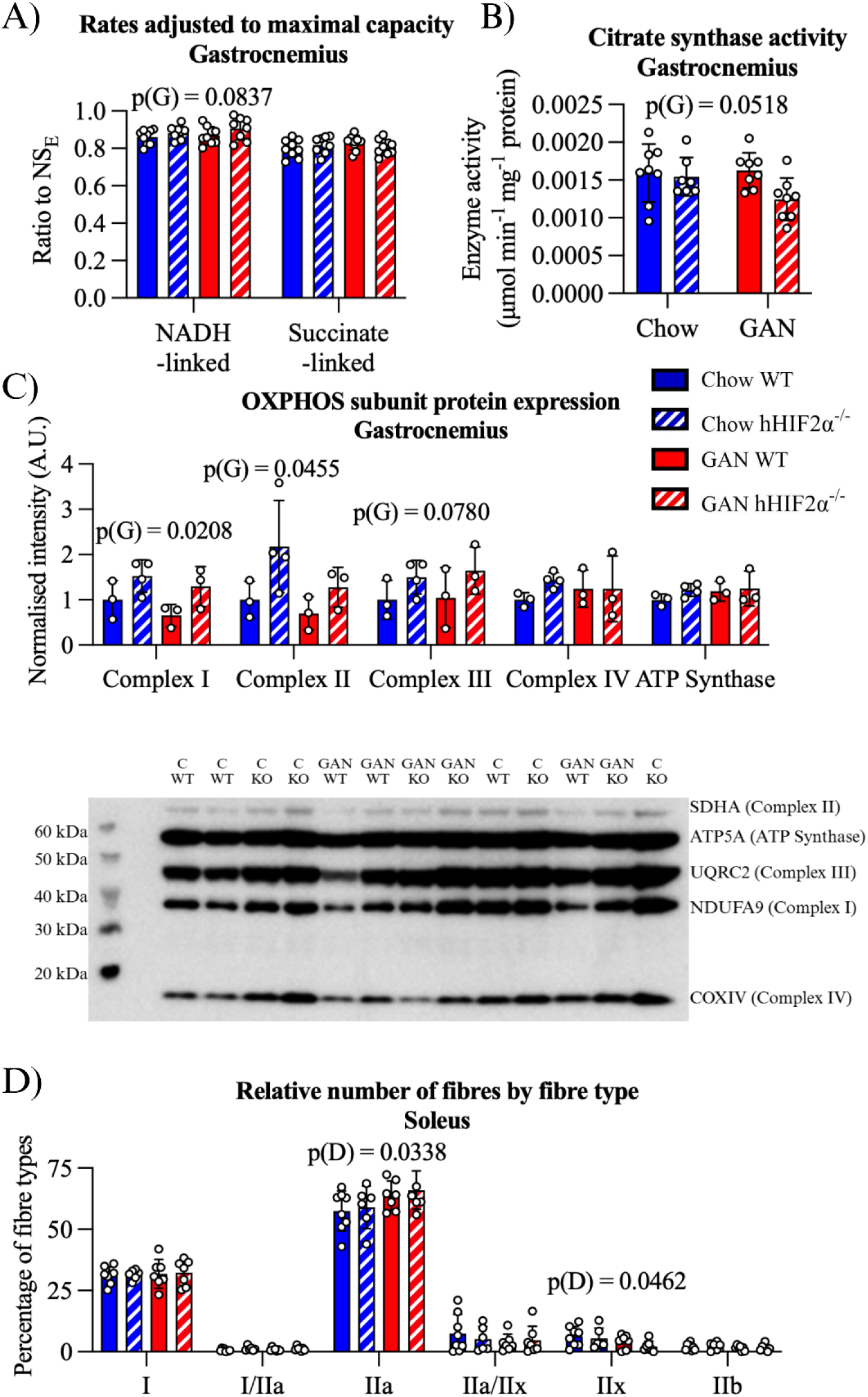
Hepatocyte-specific HIF2α deletion is associated with changes in OXPHOS protein levels in gastrocnemius muscle, while GAN-fed MASLD mice have fibre-type changes in soleus muscle. A) Respirometry of freeze-thawed gastrocnemius fibres. NADH- and Succinate-linked capacities relative to maximal (NADH- and Succinate-linked ETS combined) capacity are shown. n = 8-9 per group. B) Citrate synthase activity in gastrocnemius. n = 7-8 per group. C) OXPHOS complex subunit protein levels in gastrocnemius, normalised to Ponceau stain intensity and mean value of the wildtype chow-fed group. n = 3-4 per group. D) Relative fibre composition of soleus cross-sections. The percentage of fibre-types containing different myosin heavy chain isoforms is shown. n = 6-8 per group. See Supplementary Fig. S3D for representative images. Data are presented as mean ± SD. Results of two-way ANOVA are shown on graphs. D = Diet, G = Genotype main effect

In the primarily oxidative fibre soleus muscle, mitochondrial respiration was unaffected by genotype or diet (Supplementary Fig. S3C). Immunohistological fibre-typing (Fig. 5D) revealed a shift towards fast-oxidative type IIa fibres (p = 0.0338), away from fast glycolytic type IIx fibres in GAN-fed mice compared with chow-fed mice (p = 0.0462). Overall, gastrocnemius mitochondrial protein expression was affected by hepatocyte-specific HIF2α deletion, while GAN-feeding led to a relative shift from glycolytic to oxidative fast twitch fibres in soleus.

### Plasma lipidome is affected by GAN feeding but not hepatocyte-specific HIF2α deletion

As a potential mechanism linking changes in cardiac and skeletal muscle metabolism with the hepatocyte-specific deletion of HIF2α, we investigated the plasma lipidome. Multivariate analysis using principal component analysis (Fig. 6A) revealed separation based on diet, but surprisingly not genotype. Further investigation of the diet effect was carried out using OPLS-DA (Fig. 6B), with VIP scores of the OPLS-DA model (Fig. 6C) highlighting higher levels of the lysophosphatidylcholine LPC 18:1, several phosphatidylcholines (PC 34:1, 36:4 and 38:7), cholesterol and the cholesteryl ester CE 20:4, as well as lower levels of several triacylglycerols (TAG 52:4, 56:6, 54:5, 52:3) in GAN-fed mice relative to chow-fed mice. Similar patterns were observed when analysing overall lipid classes (Fig. 6D), with 64% lower triacylglycerol levels (p < 0.0001), 91% higher cholesteryl ester levels (p < 0.0001), 2.6-fold higher cholesterol (p < 0.0001), 23% higher sphingomyelin levels (p = 0.0137), 34% higher lysophosphatidylcholine levels (p < 0.0001) and 39% higher phosphatidylcholine levels (p < 0.0001) in GAN-fed mice than chow-fed mice. Further, the ratio of cholesteryl ester to non-esterified cholesterol was 26% lower in GAN-fed mice than chow-fed mice (p = 0.0017). No lipid class was affected by hepatocyte-specific HIF2α deletion. The sphingomyelins altered by hepatic HIF2α deletion in the liver and heart were either not detected in plasma (SM 41:1) or not affected by genotype (SM 42:2). Therefore, we found widespread alterations in the plasma lipidome due to GAN-feeding, but no clear signal that this was altered by hepatocyte-specific HIF2α deletion.

**Fig. 6:**
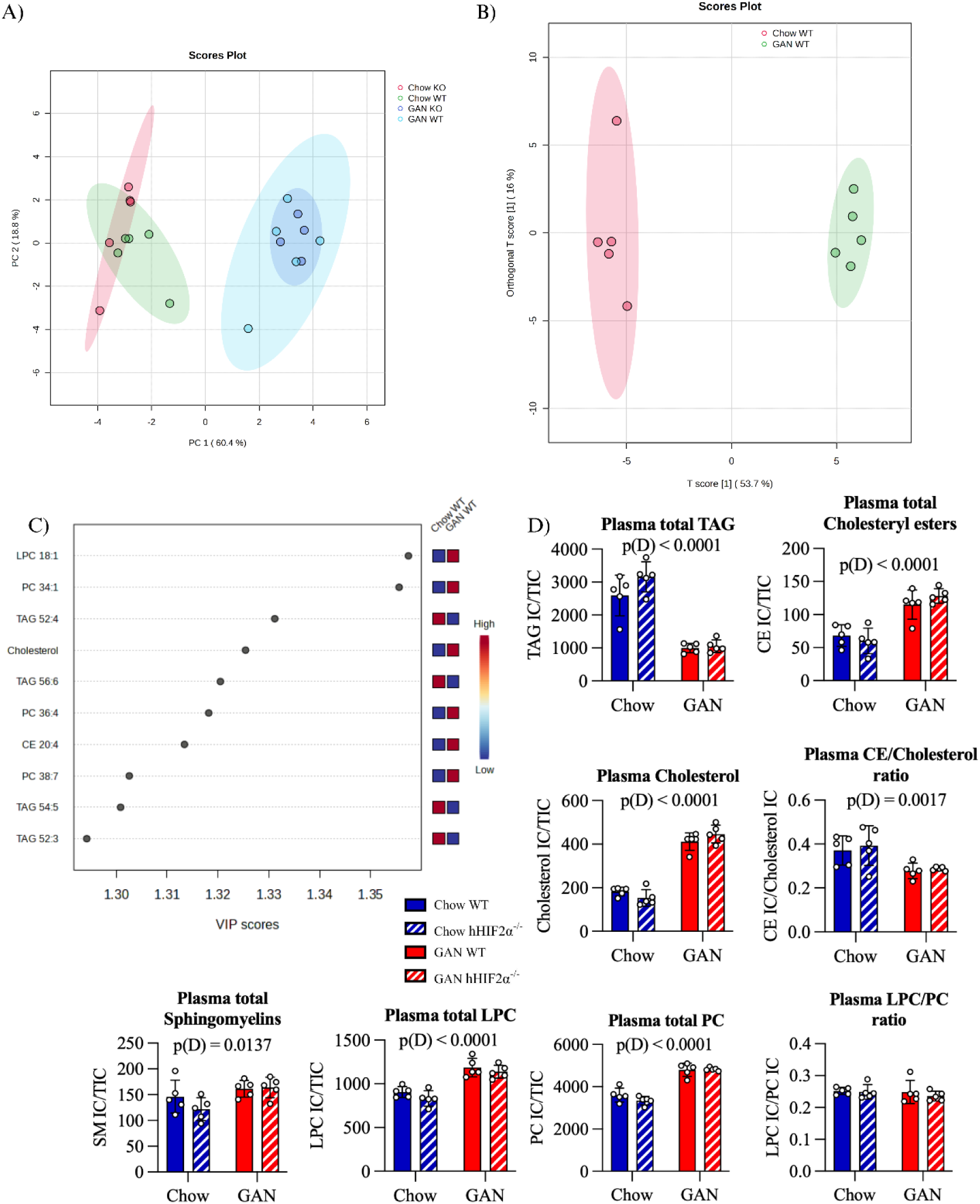
GAN feeding but not hepatocyte-specific HIF2α deletion impacts the plasma lipidome. A) Principal component analysis of plasma lipidome including all four groups. Shaded areas represent 95% confidence limits. B) Orthogonal partial least squares discriminant analysis (OPLS-DA) of diet effect in wildtype mice. Shaded areas represent 95% confidence limits. R^2^X (p1) = 0.537, R^2^Y (p1) = 0.964, Q^2^ (p1) = 0.914. For more detail on the model, see Supplemental Fig. S4. C) Top ten lipids by variable influence on projection (VIP) scores contributing to OPLS-DA model of diet effect in wildtype mice. D) Total signal for lipid classes in plasma relative to total signal of lipidome. TAG – triacylglycerol; CE – cholesteryl ester; LPC – lysophosphatidylcholine; PC – phosphatidylcholine Data presented as mean ± SD. n = 5 per group. Results of two-way ANOVA are shown on graphs. D = Diet main effect.

## Discussion

Hypoxia-inducible factor 2α (HIF2α) accumulates in the livers of patients with MASLD and has been implicated in disease progression. Here we investigated the role of hepatocyte HIF2α in driving hepatic and extrahepatic pathology using a well-validated, obese mouse model of MASLD, which replicates many aspects of the human disease^20^.

We found no apparent protective effect of hepatocyte-specific HIF2α deletion on liver pathology, adiposity or blood glucose levels, although hyperinsulinaemia was ameliorated. We had posited that HIF2α would modulate hepatic metabolism, and accordingly hepatic NADH-linked respiration was greater in hHIF2α^-/-^ mice than wildtypes. However, FAO capacity and hepatic acylcarnitine profiles were not different, despite altered expression of FAO-associated genes. Two sphingomyelin species, SM 41:1 and SM 42:2, were elevated in hHIF2α^-/-^ mice, but there were no changes in the expression of *de novo* lipogenesis-associated genes.

Despite a limited impact on liver metabolism, cardiac metabolism and function were significantly altered by hepatocyte-specific HIF2α deletion, which resulted in systolic and diastolic dysfunction. Hepatocyte HIF2α deletion also protected hearts against GAN diet-induced sympathetic dominance. Metabolically, GAN feeding resulted in myocardial lipid accumulation, particularly triacylglycerols and ceramides, and increased FAO capacity. Hepatocyte-specific HIF2α deletion alone caused myocardial accumulation of diacylglycerols and lysophosphatidylcholine, and worsened ceramide accumulation in GAN-fed mice. SM 41:1 and SM 42:2 were also elevated in hearts of hHIF2α^-/-^ mice, highlighting a possible connection between hepatic and cardiac lipid metabolism.

In skeletal muscle, there was further evidence of an extrahepatic impact of hepatic oxygen-sensing in the context of MASLD. Whole-body lean mass was lower in hHIF2α^-/-^ mice, whilst mitochondrial protein levels in gastrocnemius muscle were higher than in wildtype mice, possibly representing a compensatory response to lower lean mass, with greater relative energetic challenge experienced per unit mass of muscle.

### Hepatic HIF2α deletion does not protect against MASLD

In contrast with previous observations in the choline-deficient, lean mouse model of MASH^15^, our study revealed no protective effect of hepatic HIF2α deletion on liver pathology in GAN-fed mice, despite greater expression of hepatic HIF2α in GAN-fed wildtype mice. Expression of histidine-rich glycoprotein (*Hrgp*), determined to be central to the pathophysiological mechanism by which HIF2α contributes to MASH during choline-deficiency^15^, was not elevated in GAN-fed mice and was unaffected by hepatocyte-specific HIF2α deletion. We had posited that HIF2α-mediated regulation of hepatic lipid metabolism would influence the pathophysiology of MASLD. However, while *Cpt1a* expression was greater in GAN-fed hHIF2α^-/-^ mice than wildtype counterparts, in line with previous research^9,17^, other markers of FAO, e.g. *Hadh* expression, octanoylcarnitine-supported respiration, and the hepatic acylcarnitine pool, were unaffected by HIF2α deletion. Previous studies have indicated regulation of fatty acid synthesis by HIF2α, but with somewhat conflicting results, showing both (transiently) increased^17^, and decreased^9^ lipogenic gene expression following hepatic HIF2α overexpression. We found no effect of HIF2α on fatty acid synthase or *Srebf1* expression, and limited changes to the hepatic lipidome. Overall, our data suggest that (patho)physiological accumulation of HIF2α, in the context of long-term MASLD progression, has a more limited role in the regulation of hepatic lipid metabolism than that seen following HIF2α overexpression. Amelioration of MASLD has been reported following oral treatment with a HIF2α antagonist^33^, however this may result from inhibition of intestinal HIF2α, where deletion protects against hepatic steatosis in obesity^34^.

### Cardiac metabolic and functional changes occur in mice lacking hepatocyte HIF2α independent of MASLD

Despite the modest impact of hepatocyte HIF2α deletion on the hepatic phenotype of MASLD, we found evidence of negative extra-hepatic effects, including cardiac systolic and diastolic dysfunction. This may relate to the myocardial accumulation of putatively lipotoxic and signalling lipids, including DAGs, LPCs and ceramides^35^. Indeed, myocardial DAG accumulation following Diacylglycerol Acyltransferase 1 (DGAT1) deletion, mechanistically causes cardiac dysfunction^36^. It is unclear why DAGs accumulated in hHIF2α^-/-^ mouse hearts, but one possibility is increased activity of phospholipase C, a component of several pathways implicated in cardiac disease^37^. LPCs also accumulated in hHIF2α^-/-^ mouse hearts, and these lipids play a role in phospholipase A2 signalling in heart^38^ and have been implicated in arrhythmogenesis in cardiomyocytes^39,40^. Long-chain acyl-CoA species can reverse calmodulin-mediated inhibition of phospholipase A2^41^ and are imported into mitochondria through the action of CPT. Expression of *Cpt1b* was lower in the hearts of hHIF2α^-/-^ mice, which could lead to accumulation of cytosolic long-chain acyl-CoAs. Long-chain acylcarnitines, produced by CPT1, did accumulate in hHIF2α^-/-^ hearts following GAN-feeding, but this may suggest further FAO impairment downstream of CPT1. Similarly, ceramide accumulation, which occurred in all GAN-fed mice, was exacerbated in hHIF2α^-/-^ mice, adding to the picture of a compounding metabolic effect of the two interventions. Ceramides have been implicated in insulin resistance^42^ and apoptosis^43^, and prevention of cardiac ceramide accumulation in high-fat diet fed mice through inhibition of acid sphingomyelinase has a cardioprotective effect^44^. Lipidomic changes may therefore contribute to cardiac dysfunction in hHIF2α^-/-^ mice, which may in turn be less resilient to the additional stress imposed by the GAN diet.

While baseline cardiac function was impaired in hHIF2α^-/-^ mice, these mice were protected against GAN diet-associated cardiac sympathetic dominance. Cardiac sympathetic dominance occurs in MASLD patients^30^ and may be involved in the development of arrhythmia^45^. Hepatocyte-specific HIF2α deletion also protected against GAN-induced hyperinsulinaemia, and insulin has been shown to downregulate muscarinic receptor expression in rat atrial cardiomyocytes^46^, although no changes in *Chrm2* expression, or of the β_1_- and β_2_-adrenergic receptors were seen here.

### Mechanisms potentially linking hepatocyte HIF2α to cardiac and skeletal muscle

Lean mass was lower in hepatocyte-specific knockout mice, alongside altered skeletal muscle metabolism, although overall bodyweight was unaffected. Hepatic HIF1α regulates circulating levels of insulin-like growth factor binding protein 1 (IGFBP1)^47^ and a similar HIF2α-mediated mechanism may affect lean mass growth. IGFBP1 can also affect β-cell function^48^, potentially explaining the hypoinsulinaemic effect of hHIF2α^-/-^, which occurred without any change in blood glucose, suggesting no improvement in insulin sensitivity. Hepatic fat, rather than adipose tissue or pancreatic steatosis, is associated with increased insulin secretion^49^, implicating direct crosstalk between liver and pancreas. Whilst liver fat *per se* was unaffected by hHIF2α^-/-^, our findings may suggest that the mechanism by which hepatic steatosis affects insulin secretion is dependent on hepatic HIF2α accumulation.

Other potential mechanisms of liver-heart or liver-muscle crosstalk include hepatokines (liver-released proteins with hormone-like roles), lipid and non-lipid metabolites, and cytokines. Owing to the role of HIF2α in regulating hepatic lipid metabolism, we focussed here on lipids. Two sphingomyelins, SM 41:1 and SM 42:2, were elevated in both livers and hearts of hHIF2α^-/-^ mice. Sphingomyelins, alongside other sphingolipids including ceramides, have roles in cardiovascular disease^50^, and may mediate crosstalk between liver and heart. Plasma SM 42:2 levels were not affected by genotype however, whilst SM 41:1 was not detected in plasma. Indeed, in contrast with widespread plasma lipidomic changes induced by the GAN diet, no genotype-driven lipidomic changes were seen, potentially arguing against a direct lipid-driven crosstalk mechanism. However, it is possible that lipidomic changes were restricted to specific lipoprotein components, as seen in patients with metabolic syndrome^51^ and key signals may therefore not have been detectable through bulk analysis of the plasma lipidome.

Alternatively, mild developmental anaemia may play a role in some of the systemic findings in hHIF2α^-/-^ mice. Hepatic HIF2α plays a role in perinatal erythropoiesis, and hHIF2α^-/-^ mice are mildly anaemic during this period of development^52^. This anaemia is fully resolved a few days after birth, however perinatal anaemia may have longer term consequences on offspring health^53^. Maternal anaemia during pregnancy is associated with low birthweight^54^, which in turn is associated with lower lean mass lasting into adulthood^55^ and “catch up growth” leading to overall equal bodyweight but lower lean mass^56^. Perinatal iron restriction is also associated with cardiac dysfunction^57^ and loss of mitochondrial oxidative capacity^58^ in rat offspring. While it has not been shown whether this persists into adulthood, cardiac dysfunction following prenatal hypoxia exposure does persist^59,60^. Notably, prenatal stressors with phenotypic overlap with perinatal anaemia, e.g. fetal hypoxia secondary to placental restriction^61^, can lead to a decline in insulin levels as a result of β-cell loss later in life^62^. The hHIF2α^-/-^ mouse may therefore model perinatal anaemia, and chronic hypoxia-induced programming of health and disease^60^.

Key open questions therefore relate to the mechanisms underlying cardiac, skeletal muscle and whole-body metabolic phenotypes in hHIF2α^-/-^ mice, and whether these result from liver-released factors, or developmental programming by mild perinatal anaemia. Use of an adult-inducible hepatocyte HIF2α knockout mouse, or an anti-sense oligonucleotide approach, would avoid the confounding effect of the role of hepatic HIF2α in the developmentally-sensitive perinatal period. If extrahepatic effects are replicated in this model, investigation of the plasma proteome and metabolome may reveal alternative crosstalk mechanisms that could be useful targets in the treatment of cardiomyopathy, diabetes and sarcopenia. Alternatively, developmental programming of cardiovascular dysfunction by perinatal anaemia would itself have important implications for anaemia or chronic hypoxia during pregnancy Iron deficiency is the most common nutritional deficiency in pregnancy globally, affecting ∼50% of pregnancies^63^. In addition, more than 14 million live at extreme altitude (>3500m), thereby exposed to chronic hypobaric hypoxia, including women of reproductive age^64^. Further, chronic fetal hypoxia is the most common outcome of complicated pregnancy at sea level^65^. Understanding long-term health outcomes of the offspring in this context would therefore be of great importance. The cardiac lipidomic changes we report in hHIF2α^-/-^ mice, if replicated in a model of iron deficiency or chronic hypoxia in pregnancy, may in turn offer possible targets for intervention.

It is important to note limitations of this research. While the GAN diet aligns closely with Western dietary patterns and GAN-fed mice model human MASLD pathology^20^, plasma triglyceride levels were lower in GAN-fed mice than chow-fed mice, which differs from the human disease^66^, and this may impact some non-hepatic aspects of MASLD. Use of a hepatocyte-specific HIF2α knockout limits confounding effects of altering oxygen-sensing mechanisms in other cell-types, allowing consideration of extra-hepatic consequences of altered liver metabolism, however this does not account for any prenatal roles of hepatic HIF2α. Further, this research was carried out only in male mice. This is because pre-menopausal women^67^ and female rodents in several MASLD models^68–70^ are protected against the disease through mechanisms unlikely to involve hepatic HIF2α, leading to the decision to simplify the study design by excluding sex as a variable. Nonetheless, future investigation into findings outlined here should consider sex.

## Conclusions

Hepatocyte-specific deletion of HIF2α did not protect against MASLD in an obese mouse model of relevance to human disease, and had only limited impacts on hepatic mitochondrial and lipid metabolism. Our findings therefore diverge from reports of a protective impact of HIF2α deletion in some lean mouse models of liver disease, potentially indicating a more limited impact of this intervention on disease progression during long-term exposure to dietary stress. Instead, hepatic HIF2α deletion occurred alongside adverse systemic effects, including lower lean mass, cardiac accumulation of lipotoxic lipids, and cardiac systolic and diastolic dysfunction, although it did prevent GAN-induced hyperinsulinemia and cardiac sympathetic dominance, which may confer protection in some disease settings.

Collectively, our findings connect hepatic oxygen-sensing to extrahepatic lipid metabolism and organ function, underscoring the systemic nature of MASLD. With a substantial and growing global prevalence of MASLD and MASH, there is an urgent need for specific therapeutic interventions, and therefore a better understanding of the pathophysiology underpinning this complex condition. Our work highlights new roles for hepatic oxygen-sensing in determining systemic aspects of the condition and supports the continued pursuit of liver-targeted metabolic interventions, whilst reaffirming the importance of considering systemic physiology in the contexts of MASLD itself, and its treatment.

## Materials and Methods

### Chemicals

Unless otherwise specified, all chemicals were supplied by Sigma-Aldrich (Merck Group, Gillingham, Dorset, UK).

### Ethical Approval

All animal work was carried out in accordance with UK Home Office guidelines under the Animals in Scientific Procedures (1986) Act. All procedures involving live animals were carried out by a personal licence holder under a project licence in accordance with these regulations and received prior approval from the University of Cambridge Animal Welfare and Ethical Review Board.

### Animals

In order to obtain mice with a hepatocyte-specific deletion of *Epas1* (hHIF2α-/-), C57Bl6/J mice carrying a floxed allele of *Epas1*^71^, kindly provided by Prof. Randall Johnson, were crossed with C57Bl6/J mice expressing Cre recombinase under control of the mouse albumin promoter (strain number 003574), purchased from Jackson Laboratory (Bar Harbor, Maine, USA). Breeding was carried out by Charles River (Margate, Kent, UK). Male mice homozygous for the floxed allele and heterozygous for Cre recombinase were used as hHIF2α^-/-^ mice (n = 26), while male mice homozygous for the floxed allele but negative for Cre recombinase were used as wildtype controls (n = 27). The LoxP sites surround exon 2 of the *Epas1* allele, which encodes the basic Helix-Loop-Helix domain required for DNA binding, and recombination leads to an out-of-frame deletion of this exon ^52^. Male mice were chosen, as female rodents are relatively protected against MASLD^70^.

After weaning, mice were shipped to local facilities and allowed to acclimatise for at least one week, where they were fed standard laboratory chow diet (RM3, Special Diet Services, Essex, UK; 3.6 kcal/g, 62% kcal from carbohydrate, 11.5% kcal from fat) and housed at 21°C, 50-60% humidity, on a 12h/12h light/dark cycle. At six to nine weeks of age, mice were provided with either a high-fat, high-fructose, high-cholesterol diet (Gubra-Amylin-NASH (GAN) diet, D09100310, Research Diets Inc., New Brunswick, New Jersey, USA; 4.49 kcal/g, 40% kcal from carbohydrate (60% fructose, 40% glucose), 40% kcal from fat (43% saturated, particularly palmitate), 2% cholesterol by weight) (n = 13-14 per genotype) or continued to be fed standard laboratory chow diet (Chow, n = 13 per genotype) for 28 weeks. Mice were housed in cages of 1-4 animals, separated by diet but with mixed genotypes. Throughout the study, animals had *ad libitum* access to water, and to food except during overnight fasts before blood sampling. Fig. 1A provides an overview of the study design.

Blood sampling was carried out during week 28 after a 12-14 hour fast beginning at 9 pm. Mice were restrained in a standard acrylic mouse restraint and up to 100 µL of blood was collected from the lateral tail vein directly into a K_3_EDTA containing Microvette® (Sarstedt AG & Co. KG, Nuembrecht, North Rhine-Westphalia, Germany). Blood was immediately centrifuged (10 min, 4000 x *g*, 4°C), with plasma removed and snap frozen in liquid nitrogen. A small drop of blood was used to measure blood glucose levels using a GlucoRX HCT meter (GlucoRX Ltd, Guildford, Surrey, UK). After blood sampling, whole body composition was measured using time domain nuclear magnetic resonance (TD-NMR; LF50H Minispec, Bruker, Coventry, UK).

Three to seven days after the fasted blood sampling in week 28, mice were culled using an intraperitoneal injection of sodium pentobarbital (Euthatal; ≥ 2 g kg^-1^ bodyweight). After cessation of peripheral signs (blinking and pedal withdrawal reflexes), the chest cavity was cut open and a blood sample taken via cardiac puncture. EDTA was added at 10% weight/volume and blood centrifuged (10 min, 4000 x *g*, 4°C). Plasma was separated and snap frozen in liquid nitrogen. Fed blood glucose was also measured (GlucoRX HCT meter, GlucoRX Ltd). Heart and liver were excised and processed: Sections from each tissue were placed either into biopsy preservation solution (BIOPS: 2.77 mM CaK_2_EGTA, 7.23 mM K_2_EGTA, 6.56 mM MgCl_2_, 50 mM MES, 5.77 mM ATP, 15 mM phosphocreatine, 20 mM imidazole, 20 mM taurine, and 0.5 mM dithiothreitol, pH 7.1) for high-resolution respirometry, placed into 10% neutral buffered formalin (Q Path®, VWR Internationl Ltd., Lutterworth, UK) or frozen in optimal cutting temperature (OCT) compound (Scigen Inc, Gardena, California, USA) for histology, or snap frozen in liquid nitrogen, for lipidomic and gene expression analysis. Soleus and gastrocnemius muscles were excised and snap frozen in liquid nitrogen. Epididymal fat pads were also removed and weighed. In a sub-cohort of animals (n = 4-5 per group), no cardiac puncture was carried out and hearts were removed whole and placed in ice-cold Krebs-Henseleit bicarbonate buffer (KHB) (120 mM NaCl, 4.7 mM KCl, 1.2 mM MgSO_2_·7H_2_O, 1.2 mM KH_2_PO_4_, 25 mM NaHCO_3_, 10 mM glucose, and 1.3 mM CaCl_2_·2H_2_O) for Langendorff perfusion.

### Histology and immunohistochemistry

#### Picrosirius red staining

For picrosirius red (PSR) for fibrosis stains, and haematoxylin (Mayer’s modified, Abcam) & eosin (0.5% alcoholic) (H&E) stains, tissue was placed in 10% neutral buffered formalin for 24 hrs and then dehydrated, cleared using Histo-Clear II (National Diagnostics, Atlanta, USA) and embedded in paraffin blocks using a Leica EG1150 Embedding Center (Leica Biosystems, Milton Keynes, UK) according to standard laboratory procedures. Tissue was sectioned to 7 µm thickness using a RM2235 Microtome (Leica Biosystems), transferred to SuperFrost Plus slides (Thermo Scientific Inc.) and dried at 37°C overnight. Tissue slices were deparaffinised using xylene (Fisher Scientific, Loughborough, UK) and stained using either haematoxylin (Mayer’s modified, Abcam, UK) and eosin (0.5% alcoholic), or for PSR 0.1% Direct Red in saturated picric acid (12 g L^-1^, no. A2520; AppliChem, Darmstadt, Germany).

#### Oil red O staining

Oil red O staining of neutral lipids was carried out on 10 µm frozen sections (cut at −20°C; OTFAS-001, Bright Instruments, Luton, UK), and placed on SuperFrost Plus slides before fixation in ice-cold 10% neutral buffered formalin for 10 mins. Sections were stained in Oil red O solution (0.3% w/v in 60% isopropanol) and counterstained with haematoxylin, mounted in glycerin gelatin at 60°C and sealed using clear nail varnish. Slides were imaged using a NanoZoomer 2.0-RS (Hamamatsu, Hamamatsu City, Japan) in triplicate. Images were extracted using NDP.view2 (Hamamatsu) and analysed using ImageJ^72^. Analysis of PSR and ORO stains was carried out using a colour deconvolution approach on 20 x magnification images. For Oil red O, the percentage area stained red (with RBG values for this picked from an image) was used (with background and blue stained nuclei removed using deconvolution), while for PSR the ratio of red (collagen) to yellow-orange (non-collagenous tissue) was used (while background was removed using deconvolution) to account for large steatotic areas. For quantification of each of these colours, thresholds were adjusted based on visual inspection of several slides from each group and this threshold was then applied to all slides.

#### Skeletal muscle fibre typing

Slides were immunohistochemically stained against fibre type I, IIa, and IIx myosin heavy chain using monoclonal antibodies BA-D5, SC-71, and 6H1 (Developmental Studies Hybridoma Bank), respectively. Unstained fibres were classified as type IIb. Procedures were adapted from previously published literature^73^. Sections were air dried for 10 min and blocked in 10% normal goat serum (NGS) for 1 h, before being incubated with the primary antibody cocktail for 1h. Subsequently, sections were washed in phosphate-buffered saline (PBS) for 3 x 5 min and incubated in the dark with the secondary antibody cocktail containing Alexa Fluor IgG2b 55, IgG1 647, and IgM 488 (Life Technologies, the Netherlands) for 1 h. The PBS washing steps were repeated, and sections were incubated in PBS with 4% wheat germ agglutinin (WGA, Life Technologies, the Netherlands) for 30 min. Sections were washed and covered with Vectashield-hardset mounting medium (Vector laboratories). Images were captured at x20 magnification with a VS200 Research Slide Scanner (Olympus) using Slideview 5.0, imported to QuPath v0.5.1 and converted into RGB files. ImageJ (v.1.45g) was used to perform background correction before importing the images into MATLAB (MATLAB and Imaging Processing Toolbox 2024a, MathWorks). The SMASH toolbox (GitHub – SMASHtoolbox/Release: Version 1.0 of the SMASH Toolbox, n.d.) was used to perform segmentation and fibre type analysis in MATLAB.

### Plasma analysis

Plasma insulin levels were measured in fed and fasted plasma samples using a Rat/Mouse Insulin ELISA (Merck) according to manufacturer instructions. Insulin resistance was estimated from fasted blood glucose and plasma insulin levels using a homeostatic model (HOMA-IR) using a reference value calculated from the mean fasted insulin and fasted blood glucose levels of the chow wildtype group, rather than the human reference value of 22.5^74^. Alanine transaminase (ALT) and aspartate transaminase (AST) were measured using SimpleStep mouse ELISA kits (Abcam) according to manufacturer instructions.

### Body composition measurements

Body composition was assessed by tdNMR using a LF50H Minispec (Bruker, Coventry, UK). For this, mice were weighed using digital scales, placed in the Minispec restrainer and the relaxation times of protons within mice were measured. From this, lean mass, fat mass and free fluid mass were determined^75^, and percentage lean, fat and free fluid mass calculated. Data acquisition was carried out using Minispec software 3.0 connected to OPUS 7.0 (both Bruker).

### High-resolution respirometry

Mitochondrial respiratory capacity was measured using an Oxygraph-2k (Oroboros Instruments, Innsbruck, Austria) in permeabilised cardiac fibres as previously published^76^ or homogenised liver samples following published methodology^77^ with a modified homogenisation protocol using Potter-Elvehjem homogeniser attached to a DLH overhead stirrer (Velp Scientifica, Usumate, Italy) set to 500 rpm for 2-3 rounds of 10 s. Tissues were kept in ice-cold BIOPS after dissection. Appropriate amounts of tissue were then either permeabilised using 50 µg ml^-1^ saponin in BIOPS (for cardiac fibres) or homogenised in respiratory medium (MiR05: 0.5 mM EGTA, 20 mM HEPES, 10 mM KH_2_PO_4_, 60 mM lactobionic acid, 3 mM MgCl_2_, 110 mM sucrose, 20 mM taurine, 1 g L^-1^ bovine serum albumin).

Samples were added to Oxygraph-2k chambers containing MiR05 at 37°C followed by sequential addition of mitochondrial substrates in Substrate-(Uncoupler)-Inhibitor titration (S(U)IT) protocols as shown in Supplementary Tables 1, 2 and 3. Rates were normalised to wet weight and residual oxygen consumption after adding antimycin A was subtracted.

### Respirometry in frozen samples

Skeletal muscle respirometry was carried out in frozen samples, according to a recently developed protocol^78^. A portion of snap-frozen skeletal muscle (∼15 mg gastrocnemius, ∼6 mg soleus) was thawed in ice-cold BIOPS, then blotted dry and weighed before being transferred into 500 µL ice-cold MiR06 (MiR05 supplemented with 280 U mL^-1^ catalase). Tissue was then homogenised using a pre-cooled PBI-Shredder SG3 (Oroboros Instruments) at Level 1 for 10 s and Level 2 for 5 s. The homogenate was removed from the Shredder-tube, the tube rinsed with 50 µL MiR06 to collect the remainder of the homogenate and finally MiR06 was added to give a final concentration of 2.5 mg mL^-1^ for gastrocnemius and 1 mg mL^-1^ for soleus. Tissue homogenate was then added to Oxygraph-2k chambers maintained at 37°C.

The following protocol was employed to probe mitochondrial respiratory capacity in skeletal muscle: Initially, cytochrome *c* (10 µM) was added to reconstitute the electron transfer system^79^. Superoxide dismutase (SOD; 5 U mL^-1^) was added to convert superoxide to hydrogen peroxide and, in conjunction with catalase (in MiR06), to regenerate H_2_O and O_2_. First, malate (1 mM) and pyruvate (5 mM) were added, with no respiration expected from fragmented mitochondria. Next, NADH (4 mM) was added to stimulate the NADH-pathway via complex I (N*_E_*). To reveal maximal ETS capacity (NS*_E_*), succinate (10 mM) was added to stimulate the succinate-pathway via complex II. Then, NADH-linked respiration was inhibited with rotenone (0.5 μM) to assess succinate-pathway capacity via complex II (S*_E_*). Subsequently, antimycin A (5 μM) was used to measure residual oxygen consumption. *J*O_2_ was recorded using DatLab software 6.1 (Oroboros Instruments, Innsbruck, Austria). Respirometry measurements were performed in duplicate. Rates were normalised to homogenate concentration and residual oxygen consumption was subtracted from all values.

### Citrate synthase activity

Citrate synthase activity was assessed in triplicate in gastrocnemius samples. ∼10 mg of snap-frozen skeletal muscle tissue was added to 325 µL homogenisation buffer (20 mM HEPES, 1 mM EDTA, 0.1% Triton X-100, pH 7.2). The tissue was homogenised using a pre-chilled cooled PBI-Shredder SG3, at Level 1 for 10 sec and Level 2 for 5 sec. The homogenate was transferred to an ice-cooled Eppendorf and the PBI-Shredder SG3 tube was rinsed with another 50 µL of homogenisation buffer, which was also added to the Eppendorf. The total of 375 µL homogenate was centrifuged using a PrismR centrifuge (Labnet International) at 17,200 x *g* for 10 min at 4° C. The supernatant was removed and re-centrifuged before the final supernatant was taken and utilised for the assay.

Protein content of samples was measured using a Pierce BCA assay (Thermo Scientific Inc) according to manufacturer instructions. Each homogenate was then diluted such that 25 µL of homogenate contained 5 µg of protein. For the citrate synthase assay, each well contained 25 µL of homogenate, 269 µL CS assay buffer (20 mM Tris, 0.1 mM 5,5’-dithiobis-2-nitrobenzoic acid, 0.3 mM acetyl-CoA, pH 8.0) and 6 uL oxaloacetate (final concentration 0.5 mM). For each sample, 2 blanks were run where oxaloacetate was replaced by CS assay buffer. The reaction was measured using a FLUOstar Omega platereader (BMG Labtech) reading at 412 nm. Average slopes were calculated in Microsoft Excel.

### Open-profile lipidomics

Lipids were extracted from 10-40 mg cardiac and hepatic tissue, and 20 µL plasma using a modified Bligh and Dyer method^80^ as published previously^81^. Briefly, samples were homogenised in 600 μL methanol:chloroform (2:1), followed by addition of 200 μL chloroform and 200 μL water. After centrifugation (16,000 x*g*, 20 min), the organic and aqueous layers were separated and the extraction process repeated on the remaining protein pellet. Half of the organic layer was removed and combined with half of the aqueous layer for acylcarnitine analysis. QC samples were obtained by pooling up to 10 μL from each sample for each tissue and type of analysis. The remaining organic layer was dried under a nitrogen stream. Samples were stored at −80°C.

Analysis of lipid species was carried out using an open-profiling approach by liquid chromatography – mass spectrometry (LC-MS) using an Ultimate 3000 UPLC (LC; Dionex/Thermo Scientific, Altrincham, Cheshier, UK) and a LTQ-HPLC Orbitrap/Velos (Thermo Scientific) mass spectrometer. Dried samples were reconstituted in an appropriate volume of chloroform:methanol (2:1) (800 µL for liver, 300 µL for heart and 400 µL for plasma). 200 µL of internal standard (IS) mix (Equisplash®, supplied by Merck, supplemented with 17:0-d33 fatty acid supplied by C/D/N isotopes, Pointe-Claire, Canada) were dried down in a new glass vial, to which 980 µL isopropanol:acetonitrile:water and 20 µL reconstituted sample were added. Final concentration of IS was 0.5 µg ml^-1^. For liver, 995 µL isopropanol:acetonitrile:water and 5 µL sample were used. A blank was taken through the entire extraction and reconstitution procedure. LC-MS was carried out in both positive and negative ionisation modes with an Acquity UPLC BEH C18 Column (Waters Limited, Wilsmlow, UK) held at a temperature of 55°C and flow rate of 0.5 ml min^-1^ with a gradient run as outlined in Supplementary Table 3.

The heated electrospray ionization source was set to 375°C, desolvation temperature was 380°C, and desolvation gas flow 40 arbitrary units. All mobile phases were prepared using UPLC grade solvents and chemicals. 5 µL of sample were injected for positive ion mode and 10 µL for negative mode. QC and blank samples were included.

### Lipidomic data processing

Data were acquired using Xcalibur software (version 2.2, Thermo Scientific Inc.). Spectra were converted to .mzML files using MS Convert (Proteowizard) and processed using an R script based on the XCMS methodology. Peak identification was carried out according to the following specifications: Approximate mass resolution (FHWM) of 4 seconds; signal to noise threshold of 10; and peaks had to be present in at least 25% of samples. Peaks were annotated using an in-house automated R script, and by comparison to the LipidMaps^82^ and the Human Metabolome databases^83^. Data were normalised to total lipid ion count (TIC), log transformed and pareto scaled, and analysed using MetaboAnalyst 5.0^84^.

### Acylcarnitine analysis

Half of the organic and half of the aqueous metabolite extracts from heart and liver samples were recombined and dried down. 300 μL of methanol:water 4:1 and 3 μL of deuterated internal standard (Carnitine Splash, Avanti Research, Alabama, USA) were added. Samples were vortex mixed, sonicated (30 min) and the supernatant was centrifuged at 4⁰C at max speed for 15 minutes. The supernatant was transferred to a 96-well plate, together with a blank every 10 samples and an internal standard.

Chromatographic separations were performed using a Hypersil Gold column kept at 30⁰C on a Vanquish (Thermo Scientific), coupled with a TSQ Quantiva (Thermo Scientific) triple quadrupole mass spectrometer. The temperature of the autosampler was set to 7⁰C. The mobile phase consisted of solvent A: 0.1 % formic acid in high performance liquid chromatography (HPLC) grade water (Fisher Scientific) and solvent B: 0.1 % formic acid in acetonitrile (Supelco). When eluting the column the following gradient was used: Initial conditions were 95 % A held for 1 min followed by a linear gradient with increase of B to 100 % at 6 min, held for 2 min, with re-equilibration for 2 min giving a total run time of 10 min with a flow rate of 400 μL/min. The needle wash consisted of 10 % aqueous acetonitrile, whilst the injection volume was 5 μL. Samples were analysed by detecting the precursors of product ions using a multiple reaction monitoring (MRM) approach. The ionisation mode used by the mass spectrometer was electro spray ionisation with a capillary voltage of 3.5kV for positive ion mode. Ion source gas flows and temperatures were established using the default values provided by the Quantiva tune page software for the desired flow rate. All compound-dependent parameters were established using the Quantiva automatic optimisation protocol infusion with standards of the relevant compounds.

Acylcarnitine signal was normalised to signal intensity of a matching internal standard. Total acylcarnitine levels were then normalised to the weight of the protein pellet formed during metabolite extraction. Relative levels of free carnitine, short chain (C2-C5), medium chain (C6-C12) and long chain (C14-C20) acylcarnitine levels were calculated as fractions of the total acylcarnitine signal.

### RT-qPCR

RNA was extracted from tissue using RNeasy Plus Universal Mini kits (Qiagen) for liver samples, and RNeasy Fibrous Tissue Mini kit (Qiagen) for heart samples. Extraction was carried out according to kit instructions. 15-30 mg of tissue were used for liver and 5-15 mg for cardiac samples. RNA concentration and quality was measured using a Nanodrop ND-1000 spectrophotometer (Thermo Scientific Inc.). cDNA was synthesised using the Quantitect Reverse Transcription kit (Qiagen) following manufacturer instructions. 200 (liver) or 400 ng (heart) RNA were used. A control using a pool of all RNA samples but without reverse transcriptase and one without template RNA were included.

Assessment of mRNA transcript levels was done by RT-qPCR using the Quantinova SYBR green kit following manufacturer’s instructions in a LightCycler 480 II (Roche AG, Basel, Switzerland). Standard curves with 10-fold dilution steps starting at 20 ng of cDNA per reaction were carried out for each gene assessed to determine appropriate cDNA concentration and assess primer efficiency. 0.5 – 1 ng of cDNA were used (diluted in cDNA nuclease free water (Thermo Scientific Inc.)). Primers were either Quantitect Primer assays (Qiagen, Supplementary Table 5) or custom primers based on published sequences, purchased from Thermo Fisher Scientific or Sigma Aldrich, see Supplementary Table 6. Reverse transcriptase and template free controls were included for every gene measured. Several housekeeping genes were assessed for each tissue and the geometric mean of the most stable combination, as calculated by the BestKeeper software^85^, was used for data normalisation: *Rn18s* and *Actb*, and *Ppia* for heart and *Rn18s* and *Srsf4* for liver. The expression level of genes of interest was determined using the 2^-ΔΔCT^ method^86^, relative to housekeeping genes (referred to as BestKeeper).

### Western blotting

Protein was extracted from gastrocnemius samples using lysis buffer (200 mM Tris-HCl, 10% Triton X-100, 10 mM EGTA, 1.5 M NaCl, 25 mM Na_4_O_7_P_2_.10H_2_O, 10 mM β-glycerophosphate, 10 mM Na_3_VO_4_, pH 7.5) supplemented with cOmplete Mini protease inhibitor cocktail (Roche AG, Basel, Switzerland). Tissue was homogenised using a MagNA Lyser (Roche AG) and lysing matrix D beads (MP Biomedicals, Santa Ana, California, USA) for 3 x 20 s at a speed setting of 6000. This was followed by centrifugation (17,200 x *g*, 10 min, 4°C). The supernatant was removed to a new tube and protein concentration measured using a Pierce BCA assay (Thermo Scientific Inc.). Protein was diluted in LDS sample buffer (Thermo Scientific Inc.) with 5% β-mercaptoethanol to a concentration of 10 µg in 7.5 µL and heated to 70°C for 10 mins before being loaded onto Invitrogen NuPAGE™ 4-12%, Bis-Tris, 15 well gels (Thermo Scientific Inc.) and separated by electrophoresis in MOPS running buffer (50 mM MOPS, 50 mM Tris Base, 0.1% SDS, 1 mM EDTA, pH 7.7) using an Invitrogen XCell SureLock Mini-Cell attached to an EPS301 Electrophoresis Power Supply (Amersham Biosciences, Chicago, USA). Electrophoresis was carried out at a constant 100 V for 2 hrs 15 mins. Two samples were repeated on all gels to allow adjustment for inter-gel and blot variability. After electrophoresis, protein was transferred to a methanol activated Immobilon-P PVDF membrane at 55 mA for 1 h 10 min in transfer buffer (48 mM Tris, 39 mM glycine, 20% Methanol, 1.3 mM SDS, pH 9.2) using a semi-dry TE 77 PWR transfer system (Cytiva, Global Life Sciences Solutions, Sheffield, UK). Membranes were stained using Ponceau S (Abcam) and imaged. After destaining, membranes were blocked using 5% BSA in Tris-buffered saline (TBS) with 0.1% Tween 20 (TBS-T) for 1 h and then incubated with Total OXPHOS Antibody cocktail (1:500 in 5% BSA in TBS-T, ab110412, Abcam), washed 3 times in TBS-T, incubated with horseradish-peroxidase (HRP) conjugated secondary Rabbit anti-mouse antibody (1:10,000 in TBS-T; Number 61-6520, Invitrogen) and imaged. For imaging, membranes were incubated for 5 min with Clarity enhanced chemiluminescence (ECL) substrate (Bio-Rad Laboratories Ltd, Watford, UK). Signal was detected using Amersham Hyperfilm ECL films (Cytiva) exposed to the membrane for different lengths of time (1 s, 5 s, 10 s, 15 s, 30 s, 1 min). Films were developed using a Fuji Medical Film Processor FPM-100A (Fuji Photo Film Co., Ltd., Tokyo, Japan) and imaged. Densitometry was carried out using ImageQuantTM TL 10.2 analysis software (Cytiva, Global Life Sciences Solutions USA LLC). All densities were adjusted to total protein content per sample using Ponceau S intensity before statistical analysis.

### Isolated Langendorff heart preparation

Immediately postmortem, isolated hearts were cannulated via the aorta and perfused via the coronary arteries while lung, mediastinal, and pericardiac brown adipose tissue were dissected away. A pulmonary arteriotomy was performed. Hearts were perfused at a constant pressure of 65 mmHg. A small flexible nonelastic balloon was inserted into the left ventricle (LV) through the left atrium (LA). The balloon was filled with deionised water and attached to a rigid, deionised water-filled catheter connected to a calibrated pressure transducer (Argon Medical Devices). The volume of the balloon was set at 30 µl using a 100 µL Hamilton syringe as approximately 5–10 mmHg of left ventricular end-diastolic pressure (LVEDP) in control hearts was obtained at this balloon volume, as previously described^31^. Recirculating KHB was filtered through a 5 µm cellulose nitrate filter (Millipore) and gassed with O_2_:CO_2_ (95:5) at 37°C. After an initial stabilisation period of 15 min, basal measurements of heart rate (HR), LV systolic pressure (LVSP) and LVEDP were recorded. Baseline LV developed pressure (LVDP) was calculated as the difference between LVSP and LVEDP. The maximum (dP/dt_max_) and minimum (dP/dt_min_) first derivatives of LV pressure, as well as the contractility index and the relaxation time constant τ were calculated automatically using the M-PAQ data acquisition system (Maastricht Programmable AcQusition System).

Cardiac chronotropic and inotropic responsiveness to carbachol (carbamylcholine chloride; range: 10^-8^-10^-6^ M) and isoprenaline ((-)-Isoproterenol (+)-bitartrate salt; range: 10^-9^-10^-7^ M) was investigated. Carbachol and isoprenaline were dissolved in KHB and introduced into the heart via the compliance chamber of the Langendorff apparatus. The heart was perfused in a non-recirculating mode to avoid accumulation of carbachol or isoprenaline within the system. Recovery time (ranging from 5–15 min) was allowed between each bolus to allow HR and LVDP to stabilise to baseline values before administration of the next bolus. Sympathetic dominance ratio was calculated as the ratio of heart rate or LVDP responses to the maximum dose of isoprenaline to the response to the maximum dose of carbachol.

### Statistical analysis

Unless otherwise stated, data were analysed by two-way ANOVA (Diet*Genotype) using GraphPad Prism 10.1.1. In case of a significant Diet x Genotype interaction term, Tukey’s *post hoc* test was carried out to determine significant differences between groups. Statistical significance was set at p < 0.05. Correlation analysis with calculation of Pearson’s r and r^2^, and two-tailed p-values was also carried out using GraphPad Prism 10.1.1.

Multivariate analysis of lipidomics data were carried out using MetaboAnalyst 5.0. Principal Component Analysis (PCA) and Orthogonal Partial Least Squares Discriminant Analysis (OPLS-DA) were employed. OPLS-DA models were assessed based on their ability to fit data to the model (R^2^X and R^2^Y) and predict data based on the model (Q^2^). MetaboAnalyst 5.0 was also used to assess the hepatic lipidome using multiple two-way ANOVAs adjusted for multiple comparison using the Benjamini-Hochberg False Discovery Rate (FDR) correction.

Plots of lipidomic data were produced using MetaboAnalyst 5.0. Other plots were produced using GraphPad Prism 10.1.1. Data are presented as mean ± standard deviation. Figures were produced using BioRender.com.

## Data availability

Raw and derived metabolomics data can be accessed via the EMBL-EBI MetaboLights database^87^ at https://www.ebi.ac.uk/metabolights/MTBLS14279 or using the study identifier MTBLS14279. All data used in this manuscript are available through the University of Cambridge repository at https://doi.org/10.17863/CAM.129594^88^.

## Acknowledgments

We thank Professor Randall Johnson (Karolinska Institute, Sweden) for providing the *Epas1*^fl/fl^ mouse line used in this paper, and Sandra Pietsch for her logistical help. Further, we would like to thank the CAF staff for technical support and training, and Jenna Armstrong for experimental support.

LMWH was supported by a Wellcome Trust PhD studentship (Grant Number: 220033/Z/19/Z). This work was further supported by a Biotechnology and Biological Sciences Research Council Doctoral Training Program, Grant Number BB/M011194/1 (to PMD); 4-yr PhD studentships from the British Heart Foundation, Grant Numbers FS/17/61/33473 (to APS), FS/19/54/34889C (to AEK), FS/4yPhD/F/20/34124 (to BDT). AJM received support from the Research Councils UK, Grant Number EP/E500552/1; and Evelyn Trust, Grant Number 16/33. RMK was supported by a grant from the Research Institute Amsterdam Movement Sciences, Amsterdam. DAG was supported by the British Heart Foundation (PG/14/5/30547) and the Medical Research Council (MR/V03362X/1).

## Notes

### Competing Interest Statement

The authors have declared no competing interest.

### Summary of Updates

Update to data availability statement to include placeholder reference for all data used.

