## Supplemental material for "Hepatic HIF2α modulates extra-hepatic disease-associated phenotypes during metabolic dysfunction-associated steatotic liver disease"

### Supplementary Figures

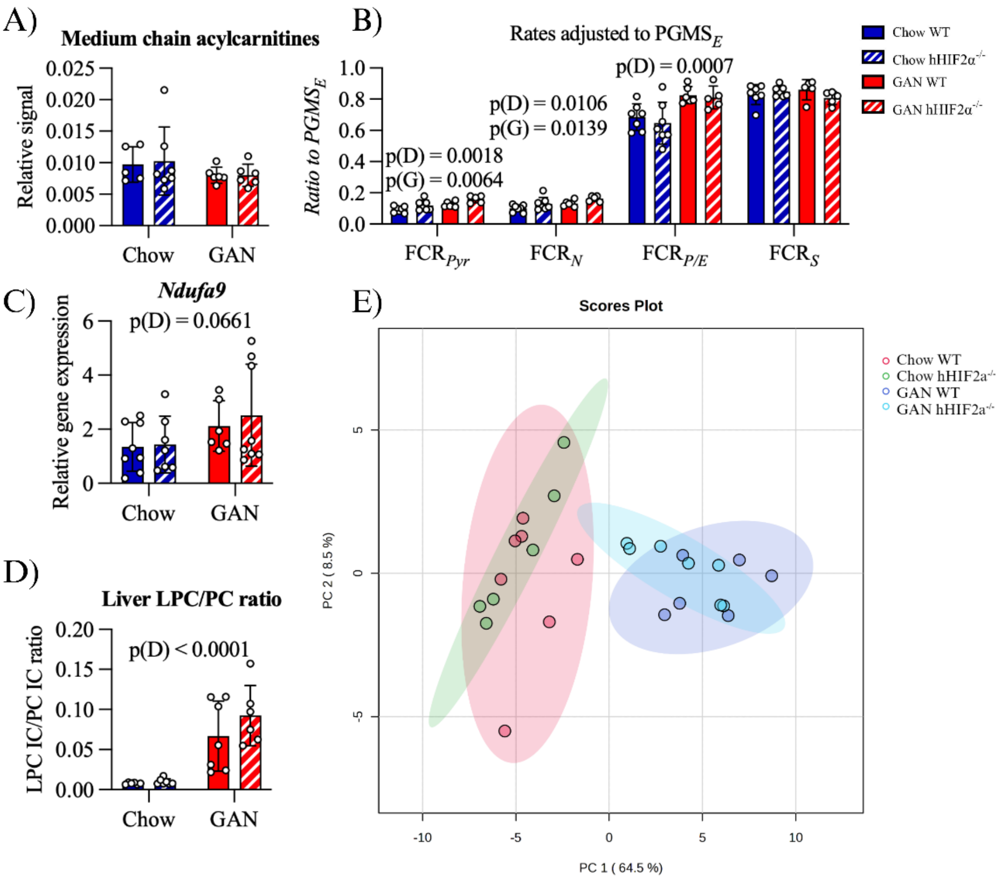


**Supplementary Fig. S1: Further hepatic metabolic data.**

A) Hepatic medium chain acylcarnitines as measured using LC-MS. n = 4-6 per group.
B) Hepatic respiratory capacities assessed using high resolution respirometry normalized to maximal electron transfer capacity (PGMS$_E$). FCR$_{Pyr}$ = PM$_P$/PGMS$_E$; FCR$_N$ = PGM$_P$/PGMS$_E$; FCR$_{P/E}$ = PGMS$_P$/PGMS$_E$; FCR$_S$ = S$_E$/PGMS$_E$.
C) Hepatic expression of the mitochondrial complex I subunit *Ndufa9* measured using RT-qPCR. n = 6-8 per group.
D) Ratio of Lysoposphatidylcholine (LPC) to phosphatidylcholine (PC) measured using LC-MS.
E) Principal component analysis of hepatic lipidome. Shaded area represents 95% confidence limits.

Data are presented as mean ± SD. Results of two-way ANOVA are shown on graphs. D = Diet, G = Genotype main effect.

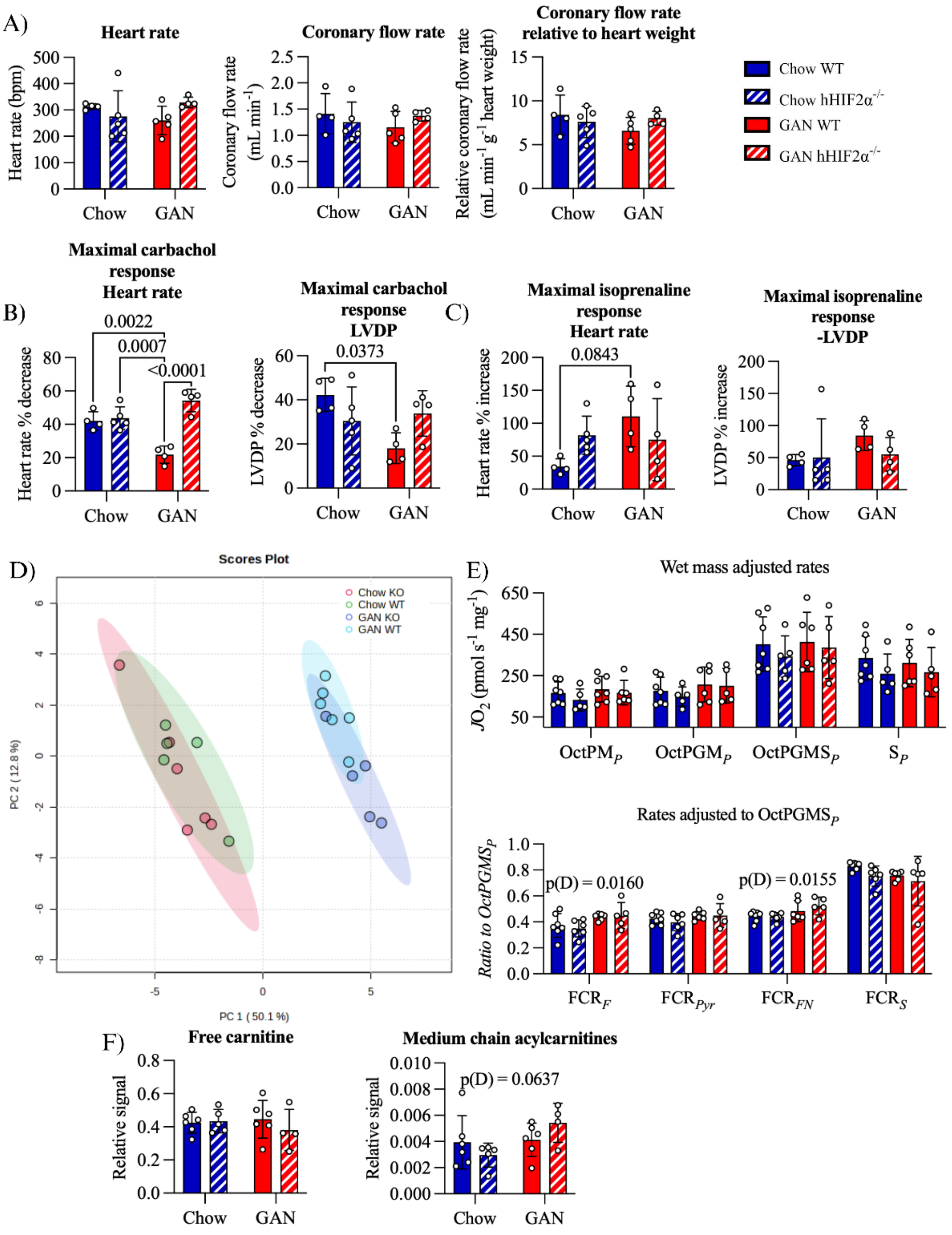

**Supplementary Fig. S2: Cardiac function in the Langendorff perfused heart and further cardiac metabolic data.**

A) Parameters of baseline cardiac function.
B) Chronotropic (heart rate) and inotropic (LVDP – left ventricular developed pressure) response to the maximal dose of the muscarinic acetylcholine receptor agonist carbachol.
C) Chronotropic and inotropic response to the β-adrenergic receptor agonist isoprenaline.
D) Principal component analysis of cardiac lipidome. Shaded area represents 95% confidence limits.
E) Cardiac respiratory capacities assessed using high resolution respirometry: Wet mass normalised capacities: Combined FAO and NADH-linked OXPHOS capacity in the presence of pyruvate, malate, octanoylcarnitine and ADP ($OctPM_P$), maximal FAO +

complex I-linked capacity after addition of glutamate (OctPGM$_P$), maximal OXPHOS capacity after addition of succinate (OctPGMS$_P$), maximal succinate (complex II)-linked OXPHOS capacity (S$_P$) after inhibition of complex I by rotenone. Flux control ratios normalized to maximal OXPHOS capacity (OctGPMS$_P$).FCR$_F$ = OctM$_P$/OctPGMS$_P$; FCR$_{Pyr}$ = OctPM$_P$/OctPGMS$_P$; FCR$_N$ = OctPGM$_P$/OctPGMS$_P$; FCR$_{P/E}$ = OctPGMS$_P$/OctPGMS$_P$; FCR$_S$ = S$_P$/OctPGMS$_P$. n = 5-7 per group.

F) Cardiac acylcarnitine pool measured using LC-MS. Free and medium chain acylcarnitines and free carnitine are expressed relative to total acylcarnitine levels. n = 4-6 per group.

Data are presented as mean ± SD. Results of two-way ANOVA are shown on graphs. In the case of a significant or near significant Diet x Genotype interaction, results of Tukey's *post hoc* test are shown.

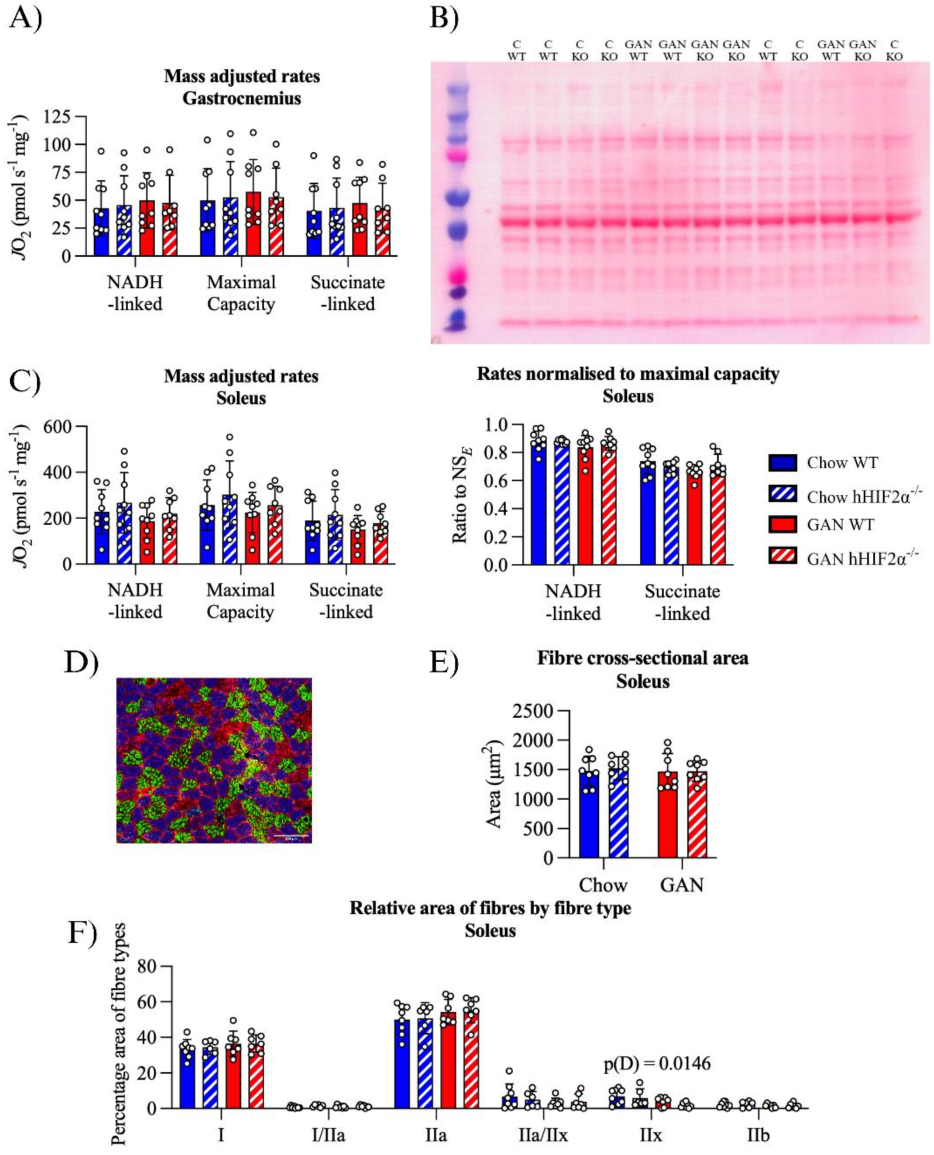


**Supplementary Fig. S3: Skeletal muscle.**

A) Respirometry of freeze-thawed gastrocnemius fibres. NADH-, maximal (i.e. NADH- and succinate-linked combined), and succinate-linked electron transfer capacities adjusted to wet mass are shown. n = 8-9 per group.

B) Ponceau stained membrane of western blot shown in Fig. 5C.
C) Respirometry of freeze-thawed soleus fibres. N-, NS- and S-linked maximal capacities adjusted to wet mass, and N- and S- linked capacities relative to maximal NS-capacity are shown. n = 8-9 per group.
D) Representative myosin heavy chain isoform-stained soleus cross-section. Scale bar represents 100 µm.
E) Total fibre cross-sectional area of soleus. n = 6-8 per group.
F) Relative area of fibres by fibre type in soleus. n = 6-8 per group.

Data are presented as mean ± SD. Results of two-way ANOVA are shown on graphs. D = Diet main effect.

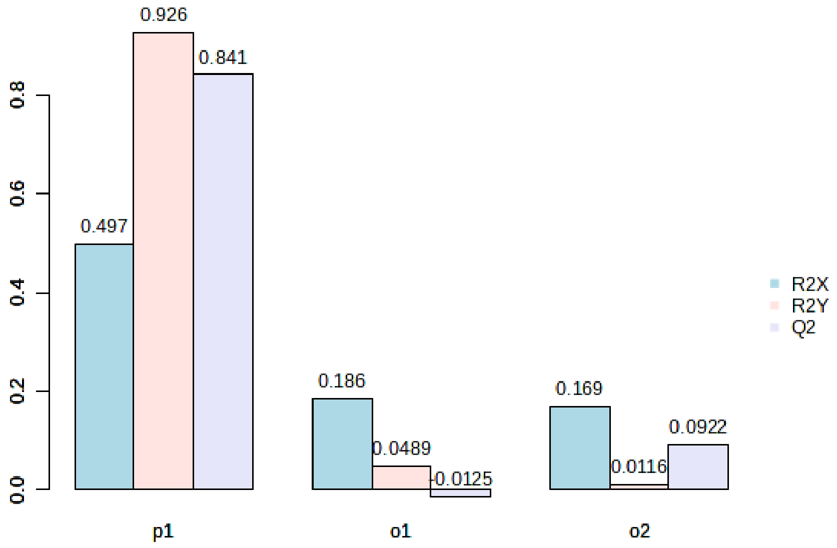


**Supplementary Fig. S4: Model performance of WT Chow – WT GAN group comparison (i.e. diet effect) on plasma lipidome.** $R^2X$, $R^2Y$ and $Q^2$ values of predictive and orthogonal components of the OPLS-DA model are shown.

### Supplementary Methods

#### Respirometry analysis

**Supplementary Table 1:** SUIT assay for liver high resolution respirometry.

| Substrate/Uncoupler /Inhibitor | Concentration (mM) | Respiratory state | Pathway probed |
|---|---|---|---|
| Malate | 1 | $PM_L$ | N-pathway (complex I) supported LEAK |
| Pyruvate | 5 | | |
| ADP | 5 | | |
| Cytochrome *c* | 0.01 | $PM_P$ | N-pathway supported OXPHOS. Cytochrome *c* assesses mitochondrial membrane integrity. |
| Glutamate | 10 | $PGM_P$ | N-pathway supported OXPHOS |
| Succinate | 10 | $PGMS_P$ | N- and S-pathway (complex II) supported OXPHOS |
| FCCP | *0.25 µM additions until maximum reached* | $PGMS_E$ | N- and S-pathway supported electron transfer system capacity |
| Rotenone | 0.0005 | $S_E$ | S-pathway supported electron transfer system capacity |
| Antimycin a | 0.0025 | Residual oxygen consumption (Rox) | Non-mitochondrial oxygen consumption |

**Supplementary Table 2:** SUIT assay for assessment of hepatic fatty acid oxidation.

| Substrate/Uncoupler /Inhibitor | Concentration (mM) | Respiratory state | Pathway probed |
|---|---|---|---|
| Malate | 1 | $OctM_L$ | Fatty acid oxidation (F-pathway) supported LEAK |
| Octanoylcarnitine | 0.5 | | |
| ADP | 5 | | |
| Cytochrome *c* | 0.01 | $OctM_P$ | F-pathway supported OXPHOS |

**Supplementary Table 3:** SUIT assay for heart high resolution respirometry.

| Substrate/Uncoupler /Inhibitor | Concentration (mM) | Respiratory state | Pathway probed |
|---|---|---|---|
| Malate | 1 | $OctM_L$ | Fatty acid oxidation (F-pathway) supported LEAK |
| Octanoylcarnitine | 0.5 | | |
| ADP | 5 | | |
| Cytochrome *c* | 0.01 | $OctM_P$ | F-pathway supported OXPHOS |
| Pyruvate | 25 | $OctPM_P$ | F- + N-pathway supported OXPHOS |
| Glutamate | 10 | $OctPGM_P$ | F- + N-pathway (complex I) supported OXPHOS |
| Succinate | 10 | $OctPGMS_P$ | F-, N- and S-pathway (complex II) supported OXPHOS |
| Rotenone | 0.0005 | $S_E$ | S-pathway supported OXPHOS |
| Antimycin a | 0.0025 | Residual oxygen consumption (Rox) | Non-mitochondrial oxygen consumption |

In addition to mass corrected rates, flux control ratios were also calculated by normalising rates to the maximal respiratory capacity measured in each assay (i.e. $OctPGMS_P$ for heart and $PGMS_E$ for liver). This mathematically removes the effect of mitochondrial content.

**LC-MS lipidomics**

Mobile phases were as follows:

Mobile phase A: 60% ACN, 40% H20 + 10 mM ammonium formate (positive)
Mobile phase B: 90% IPA, 10% ACN + 10 mM ammonium formate (positive)
Mobile phase C: 60% ACN, 40% H20 + 10 mM ammonium acetate (negative)
Mobile phase D: 90% IPA, 10% ACN + 10 mM ammonium acetate (negative)
Seal wash: 90% $H_20$, 10% IPA
Needle wash = 90% ACN, 10% IPA

**Supplementary table 4:** Gradient run used in LC-MS experiments.

| Time (min) | Mobile phase A (%) | Mobile phase B (%) |
|---|---|---|
| **0** | 60 | 40 |
| **0.8** | 57 | 43 |
| **0.9** | 50 | 50 |
| **4.8** | 46 | 54 |
| **4.9** | 30 | 70 |
| **5.8** | 19 | 81 |
| **8** | 1 | 99 |
| **8.5** | 1 | 99 |
| **8.6** | 60 | 40 |
| **10** | 60 | 40 |

**RT-qPCR**

**Supplementary Table 5:** Qiagen primers used.

| Target | NCBI Reference Sequence | GeneGlobe ID |
|---|---|---|
| ***Rn18s*** | NR_003278.3 | QT02448075 |
| ***Actb*** | NM_007393 | QT01136772 |
| ***Adrb1*** | NM_007419 | QT00258692 |
| ***Adrb2*** | NM_007420 | QT00253967 |
| ***Cers2*** | NM_029789 | QT00144025 |
| ***Cers6*** | NM_172856 | QT00137291 |
| ***Chrm2*** | NM_203491 | QT00290297 |
| ***Col1a1*** | NM_007742 | QT00162204 |
| ***Cpt1a*** | NM_013495 | QT00106820 |
| ***Cpt1b*** | NM_009948 | QT00172564 |
| ***Fasn*** | NM_007988 | QT00149240 |
| ***Hadh*** | NM_008212 | QT00147672 |
| ***Hrg*** | NM_053176 | QT00172249 |
| ***Pgc1a*** | NM_008904 | QT00156303 |
| ***Ppara*** | NM_001113418 | QT00137984 |

| | | |
|---|---|---|
| ***Ppia*** | NM_008907 | QT00247709 |
| ***Serpine1*** | NM_008871 | QT00154756 |
| ***Sgms1*** | NM_001168526 | QT00133735 |
| ***Srebf1*** | NM_011480 | QT00167055 |
| ***Srsf4*** | NM_020587 | QT01056132 |

**Supplementary Table 6:** Custom primer sequences used

| Target | Reference | Supplier | Sequence |
|---|---|---|---|
| *Epas1* deletion | Ajouaou *et al*[1] | Thermo Fisher Inc | Forward: 5′ – ACG GAG GTC TTC TAT GAG TTG GC - 3′ |
| | | | Reverse: 5′ – GTT ATC CAT TTG CTG GTC GGC - 3′ |
| *Ndufa9* | Garcia-Ruiz *et al*[2] | Sigma Aldrich | Forward: 5′ - CAT TAC TGC AGA GCC ACT - 3′ |
| | | | Reverse: 5′ - ATC AGA CGA AGG TGC ATG AT - 3′ |


1 Ajouaou, Y. *et al.* The oxygen sensor prolyl hydroxylase domain 2 regulates the in vivo suppressive capacity of regulatory T cells. *Elife* **11** (2022). https://doi.org:10.7554/eLife.70555

2 Garcia-Ruiz, I., Solis-Munoz, P., Fernandez-Moreira, D., Munoz-Yague, T. & Solis-Herruzo, J. A. Pioglitazone leads to an inactivation and disassembly of complex I of the mitochondrial respiratory chain. *BMC Biol* **11**, 88 (2013). https://doi.org:10.1186/1741-7007-11-88
